# Structure of human ferroportin reveals molecular basis of iron homeostasis

**DOI:** 10.1101/2020.03.16.993006

**Authors:** Christian B. Billesbølle, Caleigh M. Azumaya, Rachael C. Kretsch, Alexander S. Powers, Shane Gonen, Simon Schneider, Tara Arvedson, Ron O. Dror, Yifan Cheng, Aashish Manglik

## Abstract

The serum iron level in humans is tightly controlled by the action of the hormone hepcidin on the iron efflux transporter ferroportin. Hepcidin negatively regulates iron absorption and iron recycling by inducing ferroportin internalization and degradation. Aberrant ferroportin activity can lead to diseases of iron overload, like hemochromatosis, or iron limitation anemias. The molecular basis of ferroportin-mediated iron transport and regulation by hepcidin remain incompletely understood. Here, we combine cryo-electron microscopy, molecular dynamics simulations, and biochemical experiments to decipher molecular details of iron recognition and hepcidin binding to ferroportin. Iron binds to a conserved cavity in the C-domain of ferroportin, in a site unique within the broader major facilitator superfamily of transporters. We further show that hepcidin binding to ferroportin is allosterically coupled to iron binding, with an 80-fold increase in hepcidin affinity in the presence of iron. Hepcidin binds to the outward open conformation of ferroportin in a region adjacent to the iron-binding site in the C-domain. These results suggest a new model for hepcidin regulation of ferroportin, where only iron loaded ferroportin molecules are targeted for degradation. More broadly, our structural and functional insights are likely to enable more targeted manipulation of the hepcidin-ferroportin axis in disorders of iron homeostasis.

## Introduction

Iron is essential for life. Complexed to heme, iron enables oxygen transport and cellular respiration. As a cofactor for many proteins, iron coordinates redox chemistry by alternating between ferrous (Fe^2+^) and ferric (Fe^3+^) oxidation states. Despite this central role in biology, free ferrous iron is toxic. In excess, iron can catalyze the production of free radicals, leading to cellular damage. Iron levels are therefore tightly controlled, both at the cellular and organism level

In mammals, iron levels are regulated by the action of hepcidin, a peptide hormone, on ferroportin (FPN), the only known iron efflux transporter^1–3^. FPN mediates absorbance of dietary iron by transport of ferrous iron across the basolateral surface of intestinal enterocytes. FPN also mediates iron recycling from hepatocytes and macrophages^4^. Iron efflux by FPN is controlled by the amount of transporter located at the cellular surface. FPN synthesis is transcriptionally regulated by cellular hypoxia, iron and heme concentrations, and inflammatory signaling^5^. In settings of elevated serum iron levels, liver-derived hepcidin levels increase, and this hepcidinnegatively regulates cell surface FPN by both acutely blocking iron transport^6^, and by inducing FPN ubiquitination, internalization, and degradation^7–10^. Hepcidin activity decreases serum iron levels by suppressing FPN-mediated dietary iron absorption and release of iron from cellular stores.

Iron disorders in humans can result from dysregulation of hepcidin and FPN, reflecting the central role of the hepcidin-FPN axis in iron homeostasis. Deficits in hepcidin-mediated regulation of ferroportin, often due to hereditary hemochromatoses, lead to iron overload and widespread tissue affecting the liver, pancreas, and joints^11–13^. By contrast, inappropriate elevation of hepcidin levels yields iron-restricted anemia^14,15^. Although several approaches to restore aberrant FPN function have been evaluated in clinical trials^16–18^, none have thus far succeeded.

The molecular mechanism of FPN regulation by hepcidin remains incompletely defined at the atomic level. A confluence of human genetics studies and structure-function evaluations have identified key regions of FPN important in hepcidin regulation^6,19–22^. A key recent advance was determination of the X-ray crystal structure of a divalent metal transporter from the bacterium *Bdellovibrio bacteriovorus* (bbFPN) with 40% similarity to human FPN^23,24^, which revealed a unique architecture among the broader major facilitator superfamily (MFS) of membrane transporters. Although bbFPN is predicted to share structural features with human FPN, the precise mechanisms of iron coordination likely differ, and bbFPN is not regulated by hepcidin.

To understand how FPN transports iron, and how this process is regulated by hepcidin, we used a combination of structural biology, molecular dynamics simulations, and *in vitro* biochemical assays. These studies reveal the molecular recognition of iron and hepcidin by FPN and suggest a new regulatory mechanism enabling hepcidin to selectively target actively transporting ferroportin molecules for degradation.

### Structure of human FPN in a lipid nanodisc

We screened the antigen-binding fragments (Fabs) of antibodies previously raised against FPN^25^ to identify one that could be used as a fiducial mark to guide image alignment of a small membrane protein embedded in lipid nanodisc for structure determination by single particle cryogenic electron-microscopy (cryo-EM), a strategy we proposed many years ago^26^. Among the many Fabs that bound purified ferroportin (Supplementary Fig. 1), a single clone, Fab45D8, yielded interpretable class averages in negative stain EM and was selected to facilitate cryo-EM structure determination. Unlike many antibodies and antibody fragments targeting FPN, Fab45D8 was previously determined to be non-competitive with hepcidin and, on its own, did not induce FPN internalization^25^. Indeed, in nanodisc-reconstituted preparations of FPN, Fab45D8 neither altered iron affinity, nor did it change the binding properties of hepcidin (Supplementary Fig. 2).

We obtained a 3.2 Å cryo-EM density map of nanodisc-reconstituted FPN bound to Fab45D8 (Fig. 1a & Supplementary Fig. 3). This map enabled building of an atomic model of FPN residues 17-238, 291-393 and 451-547 which comprise previously identified regions important for iron transport and hepcidin binding^1,6^, and a portion of the intracellular loop 3 (ICL3) important in hepcidin-induced FPN internalization^8–10^(Supplementary Fig. 4 & Supplementary Table 1). The entire FPN extracellular loop 5 (ECL5) remains unresolved, likely due to significant conformational flexibility. To enable modeling of Fab45D8, we obtained its X-ray crystal structure at 2.1 Å (Supplementary Fig. 5 and Supplementary Table 2)

**Figure 1.**
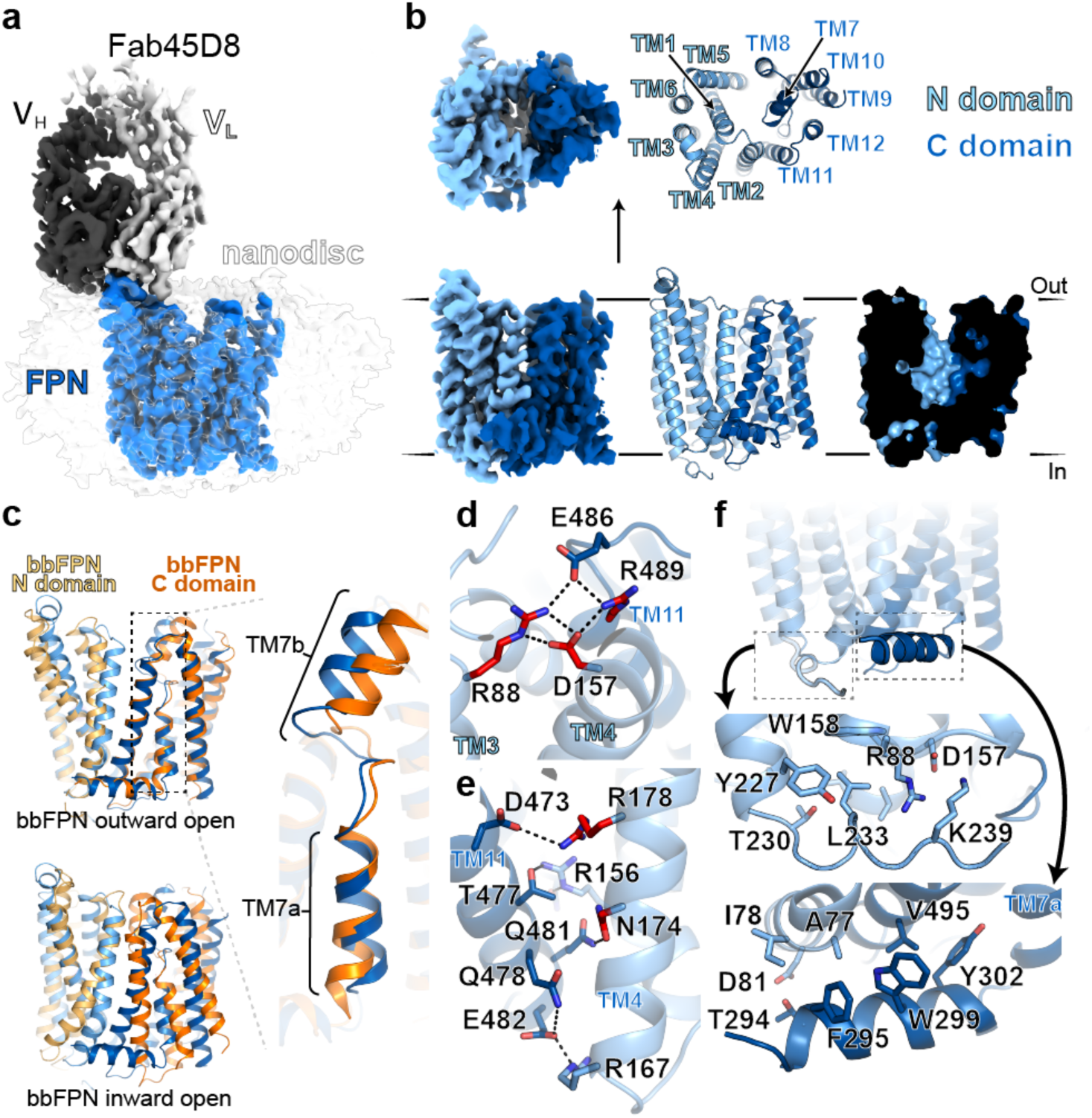
Structure of human ferroportin and comparison to bbFPN. **a**, Cryo-EM map of FPN-Fab45D8 complex in lipid nanodisc. **b**, Cryo-EM density and ribbon diagram of FPN reveals 12 transmembrane helices. The N- and C-domains are colored in different shades of blue. Cutaway surface view (right) shows outward open conformation. **c**, Human FPN aligned to the outward-open (PDB: 5AYN) and inward-open (PDB: 5AYO) conformations of bbFPN. Inset shows unique architecture of TM7 shared between human FPN and bbFPN. **d**, Intracellular gating residues are shown as sticks. Residues in red are known FPN loss-of-function mutations **e**, TM11 and TM4 form an extensive network of interactions, further stabilizing the inward-open state. Residues highlighted in red are known loss-of-function mutations that lead to ferroportin disease. **f**, Closeup views showing multiple contacts between the transmembrane helices and residues within intracellular loop 3.

Our structure reveals a monomeric FPN bound to a single Fab45D8 molecule, which recognizes a short alpha helical segment in extracellular loop 2 (ECL2) (Fig. 1a and Supplementary Fig. 5). Similar to other MFS transporters, FPN contains twelve transmembrane (TM) helices arranged in two domains. Both the N-terminal and C-terminal domains are composed of six helices, with a large central cavity that in our structure is open to the extracellular side and closed intracellularly (Fig. 1b). Ferroportin shares significant structural similarity with the bacterial bbFPN transporter, with an overall root mean squared deviation (RMSD) of 2.0 Å when compared to the outward-open conformation of bbFPN (Fig. 1c). The overall backbone conservation is even higher within the isolated C-terminal domain (RMSD 1.4 Å). Unlike most other MFS transporters, the alpha helix of FPN TM7 is interrupted by a short non-helical stretch of six residues. This unique feature, previously posited to be important in iron binding^24^, is shared between human FPN and bbFPN.

Several interacting residues define an intracellular gate that keeps the N- and C-domains of ferroportin in an outward open conformation. Similar to a previously observed interaction network in bbFPN^23^, R489 in TM11 of the C-domain forms an ionic interaction with D157 in TM4 of the N-domain (Fig. 1d). This interaction is further supported by an extended ionic and hydrogen-bonding network including residues E486 (TM11) and R88 (TM3). In human FPN, an additional cluster of ionic and hydrogen bonding interactions between TM4 in the N-domain and TM11 in the C-domain further stabilizes the inward open conformation (Fig. 1e). Mutation of several residues within the intracellular gate leads to FPN loss of function in ferroportin disease, highlighting the importance of the gate in coordinating iron efflux^19,27^.

Hepcidin regulates ferroportin by causing ubiquitination of lysine residues in intracellular loop 3 (ICL3)^9,10^. Among these, K240 is critical for hepcidin-induced FPN internalization and degradation. Our structure of ferroportin does not resolve residues 239-290 of ICL3, precluding a structural understanding of how hepcidin regulates the conformation of K240. However, the resolved region provides clues into the role of ICL3 in FPN function. The N terminal portion of ICL3 (residues 230-238) forms interactions with the N domain (Fig. 1f). Notably, K236 makes an ionic interaction with the intracellular gate residue D157. The C terminal portion of ICL3 (residues 291-304) forms an amphipathic helix that makes a number of contacts with both the N and C domains (Fig. 1f). In both cases, the resolved regions of ICL3 are primed to sense the conformation of the transporter as it shuttles iron and binds hepcidin. These regions may therefore serve as important conduits linking the conformation of hepcidin binding on the extracellular side to K240 conformation on the intracellular side.

### An iron binding site in the C domain

Ferroportin has previously been proposed to possess two distinct sites capable of binding divalent cations. Although initial crystallographic studies suggested that iron primarily binds in a cavity within the N domain of the bbFPN transporter^23^, further mutagenesis studies found a critical divalent cation binding site within the C domain^24^. Structural elucidation of the FPN iron binding site, however, remains elusive. Previous studies on bbFPN either used supraphysiological concentrations of iron or found Ni^2+^ bound as an EDTA complex.

We observed a density for a potential ion within the C domain of human FPN located adjacent to H507 in TM11 (Fig. 2a). Although two anionic residues, D325 in TM7b and D504 in TM11, may serve as additional coordinating residues for a divalent cation in this region, the side chain density of D325 is poorly resolved, consistent with known challenges in resolving carboxylates in cryo-EM maps^28^. We are unable to confidently assign the origin of this density, as we collected cryo-EM data in the absence of exogenous iron or other divalent metals and the local resolution precludes clear assessment of coordination geometry. To assess whether this site is capable of binding iron, we performed all-atom molecular dynamics simulations of FPN in a hydrated lipid bilayer. We first performed six simulations, with Fe^2+^ ions initially positioned randomly in bulk solvent. In all six independent simulations, Fe^2+^ ions bound spontaneously to the hypothesized iron binding site, localizing to the unwound region of TM7 near residues D325, D504, and H507 within hundreds of nanoseconds of simulation time (Fig. 2b,c and Supplementary Fig. 6).

**Figure 2.**
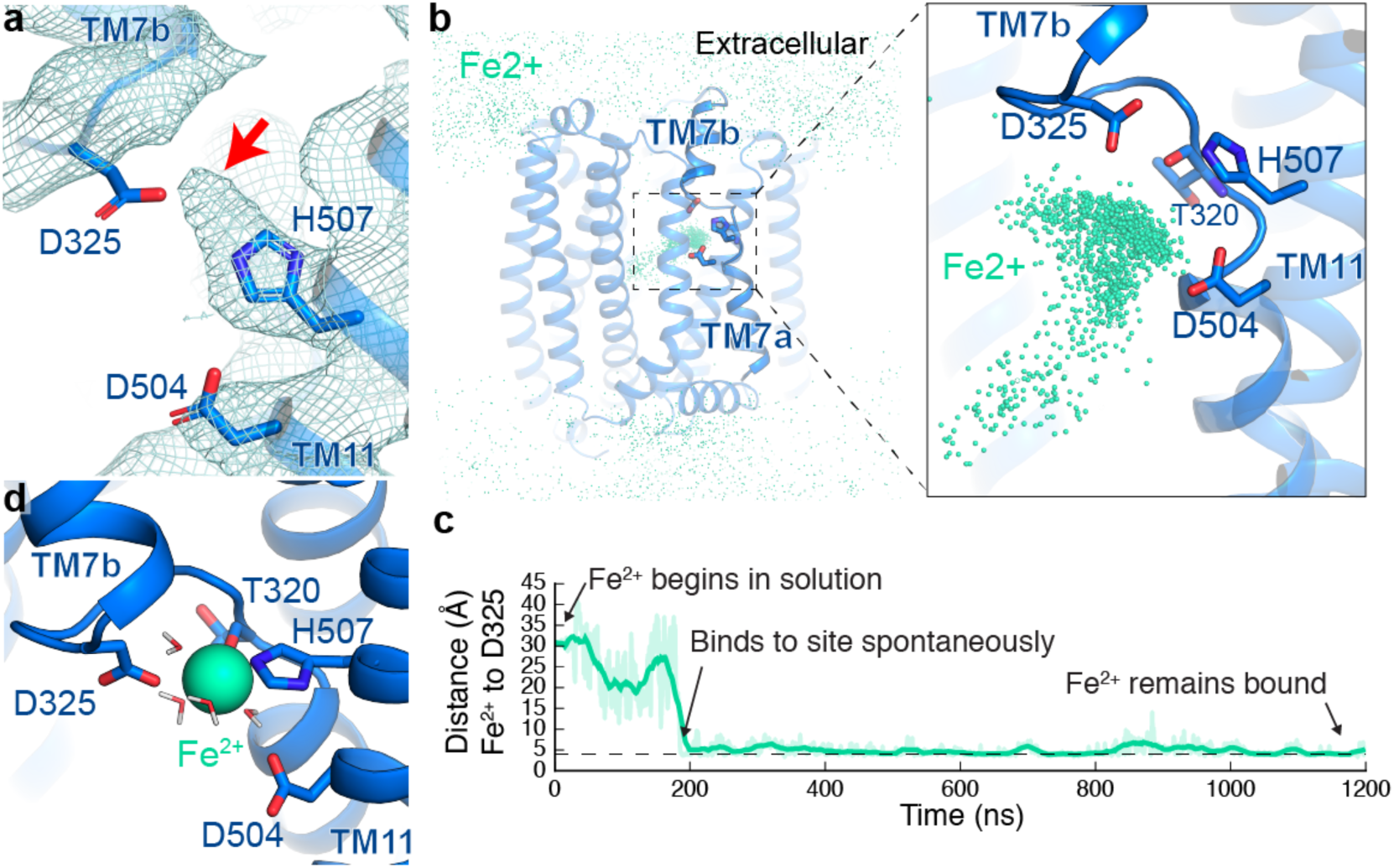
Iron binds within the C domain of FPN. **a**, Closeup view of cryo-EM density near H507, D325, and D504. The unaccounted extra density is potentially a coordinated ion. **b**, In molecular dynamics simulations with Fe^2+^ initially positioned randomly in bulk water surrounding FPN, the Fe^2+^ ions spontaneously bind to a region near H507, D325, and D504. The aggregated position of Fe^2+^ ions from six simulations, each 2 μs in length, is shown superimposed with FPN. **c**, In one representative simulation, an Fe^2+^ ion spontaneously binds within 200 ns and remains localized at this site for more than 1000 ns. Distance shown is from the ion to the nearest oxygen atom of the D325 side chain. Thick trace represents a 15-ns sliding mean and thin traces represent unsmoothed values**. d**, When Fe^2+^ ions were initially placed at the putative C domain site, they remained persistently bound. A representative frame from these simulations shows that the Fe^2+^ ion interacts with side chains of H507, D325, and D504 and the backbone carbonyl of T320.

To sample iron binding more exhaustively, we performed additional simulations where Fe^2+^ was initially placed in the proposed binding site. Here, Fe^2+^ remains stably bound; it occasionally moves closer to TM1 to interact with D39 but quickly returns to its primary location (Supplementary Fig. 6). Bound Fe^2+^ formed persistent interactions with D325, H507, D504, and the backbone carbonyl of T320 (Fig. 2d), with most frequent coordination by the side chains of D325 and H507 (Supplementary Fig. 6). In some simulation snapshots, interactions between FPN and Fe^2+^ could also be mediated by waters that coordinated the ion with an octahedral geometry. Coulombic interactions are also likely important to the observed localization at this site; the high concentration of negative charge at this site due to the two aspartates likely pulls the Fe^2+^ ion into the site quickly and keeps it from escaping. The geometry of side chains at this site suggests that only two or three side chains can directly coordinate the Fe^2+^ ion simultaneously; this is distinct from iron binding sites in other proteins such as oxygenases^29^ and may reflect the weaker ion binding needed for efficient transport.

### Hepcidin is allosterically coupled to iron binding

Hepcidin has been proposed to bind the outward-open conformation of ferroportin by making important interactions with key residues located within TM7. Indeed, the C326S mutation in TM7b results in iron overload (hemochromatosis type 4B) characterized by a lack of hepcidin binding and ferroportin degradation^30,31^ and elevated hepcidin levels. In our attempts to reconstitute a hepcidin-ferroportin complex, we noted that hepcidin displays a relatively low affinity for purified ferroportin. A fluorescently tagged version of hepcidin (Rhodamine green-hepcidin, RhoG-Hep) bound to nanodisc-reconstituted ferroportin with an apparent K_D_ of 210 nM (pK_D_ = -6.67± 0.02) (Fig. 3a). As this affinity exceeds the reference range of hepcidin concentrations found in healthy adults, which range from ∼1-30 nM^32^, we speculated that another layer of regulation may control hepcidin activity at ferroportin.

**Figure 3.**
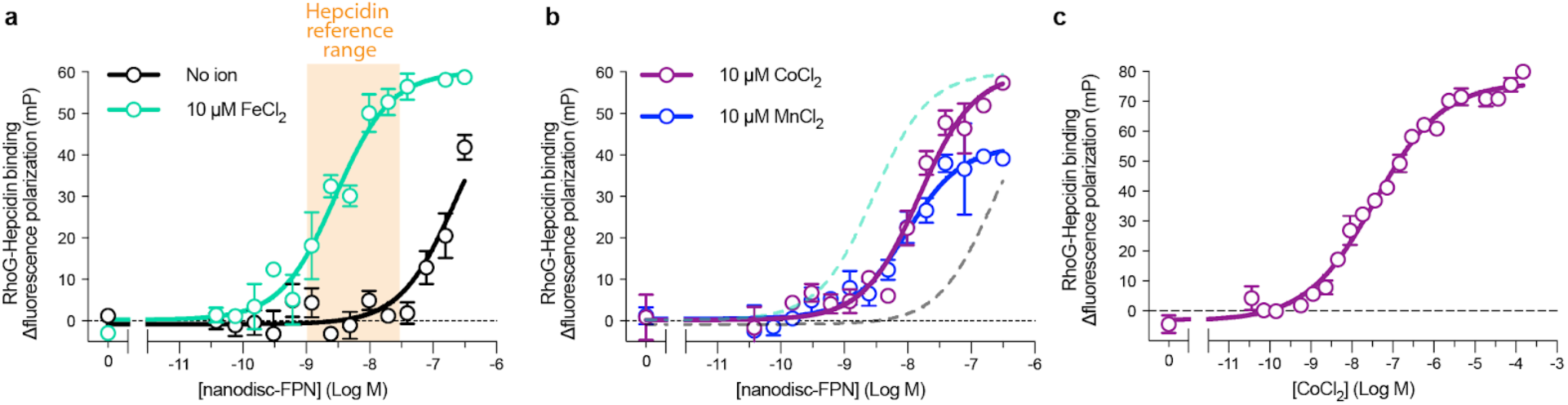
Hepcidin is allosterically coupled to iron binding. **a**, Fluorescence polarization increase in rhodamine G-labeled hepcidin (RhoG-hepcidin) as nanodisc-reconstituted FPN is titrated with a K_D_ of 210 nM. Addition of 10 µM FeCl_2_ increases the affinity of hepcidin to 2.5 nM. Hepcidin concentration range in healthy human adults is shown in orange. **b**, The transition metals Co^2+^ and Mn^2+^ similar in size to Fe^2+^ also increase RhoG-hepcidin affinity at FPN, with respective affinity of 7.7 nM (pK_D_ = -8.11 ± 0.16) and 7.8 nM (pK_D_ = -8.10 ± 0.06) **c**, Potentiation of RhoG-hepcidin binding by cobalt is saturable, with EC_50_ of 36 nM. Shown here is a single representative experiment from *n* = 3 distinct experiments. All values are reported as mean ± s.e.m. Error bars represent s.e.m.

Given the close proximity of the iron binding site to the putative hepcidin site in TM7b, we hypothesized that iron may regulate hepcidin affinity. In the presence of 10 µM FeCl_2_, we observed a significantly increased affinity of 2.5 nM (pK_D_ = -8.61 ± 0.21) between hepcidin and ferroportin (Fig. 3a). Though iron is the preferred substrate, other transition metals, including Co^2+^ and Zn^2+^ are also transported by FPN^33^. In our hands, cobalt (Co^2+^) and manganese (Mn^2+^) also increased hepcidin affinity (Fig. 3b), despite conflicting reports on the ability of FPN to transport Mn^2+33–36^. Consistent with an allosteric effect on hepcidin affinity, the increase in hepcidin affinity by Co^2+^, is saturable (Fig. 3c).

We turned again to molecular dynamics simulations to provide a structural rationale for the allosteric coupling between hepcidin and iron binding. Simulations of ferroportin without iron present revealed movements of the entire backbone of TM7b, with significant fluctuations of the key iron coordinating residue D325 (Fig. 4a-e). By contrast, both D325 and TM7b are less mobile in the presence of bound iron. In iron-bound simulation, the extracellular end of TM7b also tilts toward TM1, altering the average width of the extracellular cavity (Figure 4a,c). We therefore speculate that iron-dependent coordination of D325, and thereby stabilization of TM7b, provides a critical interface for high affinity hepcidin binding. Hepcidin residues may also directly coordinate iron.

**Figure 4.**
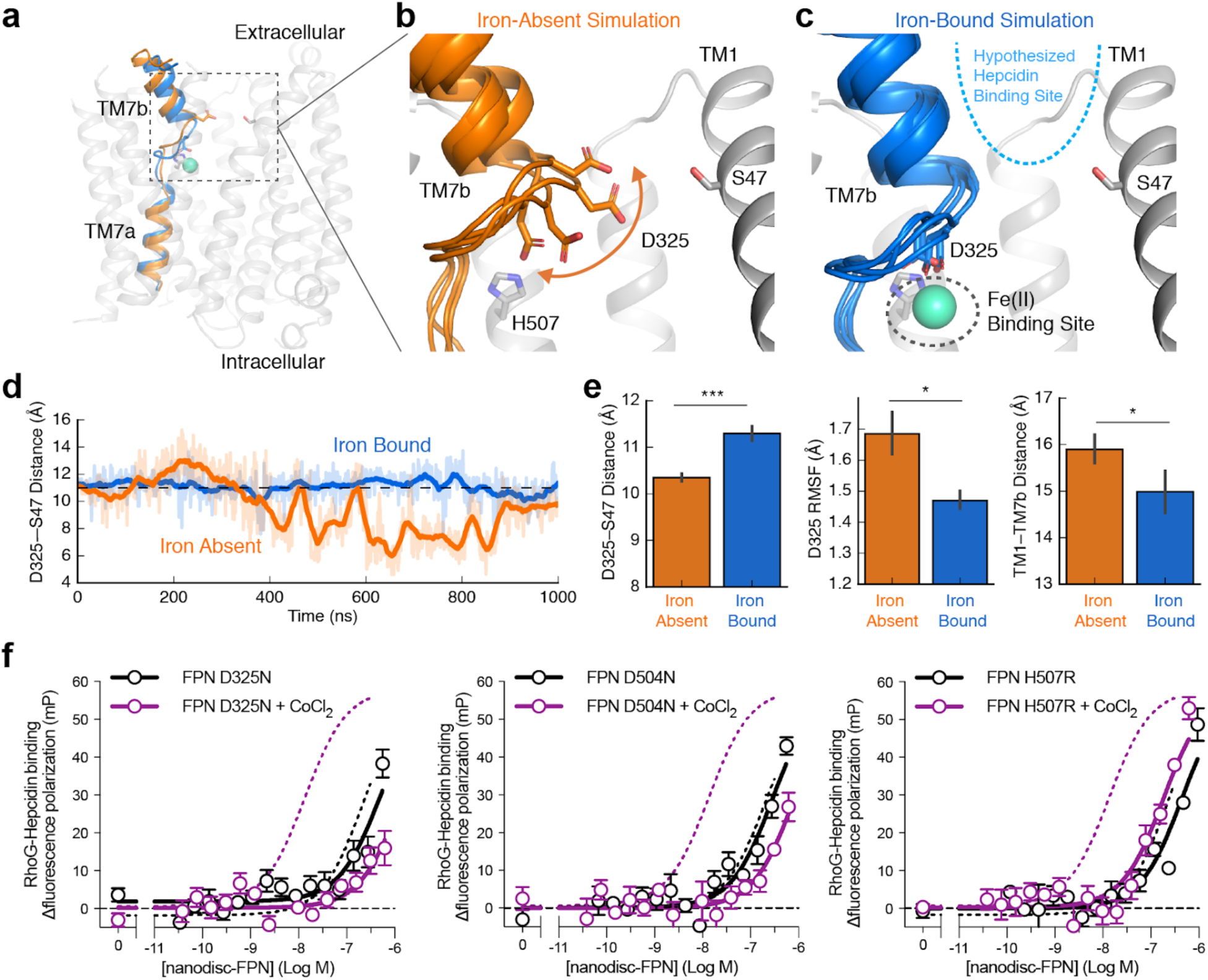
Molecular mechanism for hepcidin-iron allostery. **a**, Representative conformations of TM7 from simulations with iron bound (blue) or absent (orange). **b**, In the absence of bound iron, D325 is mobile and can move into the cavity between TM7b and TM1. The top of TM7b can also tilt away from TM1. Four representative frames from simulation are overlaid. **c**, In simulations with Fe^2+^ bound, interaction with the ion restricts the mobility of D325 and, in turn, TM7b. **d**, Iron-bound and iron-absent simulations show differences in the dynamics and position of D325, as measured by the distance between D325 Cγ and S47 Cβ. **e**, Comparison of conformation and dynamics with and without iron bound. With iron bound, D325 moves away from TM1 into the iron binding site (left), the root-mean-square fluctuation (RMSF) of D325 decreases (middle), and the extracellular end of TM7b moves closer to TM1 (right). For these comparisons, 6 simulations for each condition were used, each 2.0 μs in length. Error bars are s.e.m. and p-values were calculated using Mann-Whitney U test (* p<0.05, *** p<0.001). **f**, RhoG-hepcidin binding to FPN mutants shows micromolar binding to FPN in the absence of CoCl_2_. Compared to wild-type FPN (dotted lines), mutations at the proposed iron binding site (D325N, D504N, and H507R) lead to minimal increase in hepcidin affinity in the presence of 50 µM CoCl_2_. Each datapoint represents replicates with *n* = 3; error bars are s.e.m.

The simulations presented here, and prior mutagenesis studies, predict that TM7b and TM11 are important for iron binding. Human disease mutations suggest that the hepcidin binding site is located in a similar region of FPN. We tested our proposed iron binding site and its effect on hepcidin affinity by measuring RhoG-hepcidin binding for designed mutations lining either the iron-binding site or important hepcidin contacts. To deconvolute direct effects of mutations on hepcidin affinity, we first tested whether mutation of the putative iron binding residues D325, D504 and H507 affects RhoG-hepcidin binding. Compared to wild-type FPN, D325N, D504N, and H507R show similar micromolar binding to RhoG-hepcidin (Fig. 4f), suggesting they are not directly associated with hepcidin binding. In stark contrast to the 30-fold increase in hepcidin affinity observed for wild-type FPN in the presence of Co^2+^, the D325N, D504N, and H507R mutations show almost no gain in hepcidin affinity under similar conditions (Fig. 4f). These studies highlight the critical role of the C domain iron site in potent hepcidin binding to ferroportin.

### Visualization of hepcidin binding to ferroportin

To understand how hepcidin binds and regulates FPN, we additionally obtained cryo-EM data for nanodisc-reconstituted ferroportin in the presence of Co^2+^ and hepcidin (Supplementary Fig. 7 and Supplementary Table 1). Despite similar data collection and processing strategies as for apo ferroportin, we were unable to obtain a high resolution single particle cryo-EM map of the ferroportin-hepcidin complex (data not shown). Using UltrAufoil gold grids allowed us to overcome previous difficulties in maintaining Fab45D8 binding to FPN in vitreous ice without continuous support. This strategy enabled reconstruction of a map at 6.6 Å resolution, which showed density in the extracellular-facing FPN cavity proposed to be the site of hepcidin binding^1^. We therefore calculated difference maps between FPN analyzed in the presence and absence of Co^2+^:hepcidin after filtering both maps to 10 Å resolution.

The difference map subtracting apo FPN density from the FPN:Co^2+^:hepcidin dataset revealed a strong density located within the central cavity of FPN (Fig. 5a,b). Human FPN mutations leading to hepcidin resistance surround this difference density (Fig. 5c). Additionally, hepcidin from a previously determined X-ray crystal structure of the hormone^21^, docks into the observed difference density with the disulfide-rich region facing extracellularly and the N-terminus inserting deep into the transmembrane cavity. This binding mode is consistent with structure-activity relationship studies of hepcidin that demonstrate the functional effect of hepcidin on FPN can be recapitulated by the first nine residues of the hormone^37,38^. While the resolution of the difference map precludes confident modeling of hepcidin, this speculative binding mode places the N-terminus of hepcidin in close contact with FPN residues H507, D325, and D504 (Fig. 5c). The observed potentiation of hepcidin affinity by Fe^2+^ or Co^2+^ may therefore be driven by a direct interaction between the hepcidin N-terminus and the metal substrate.

**Figure 5.**
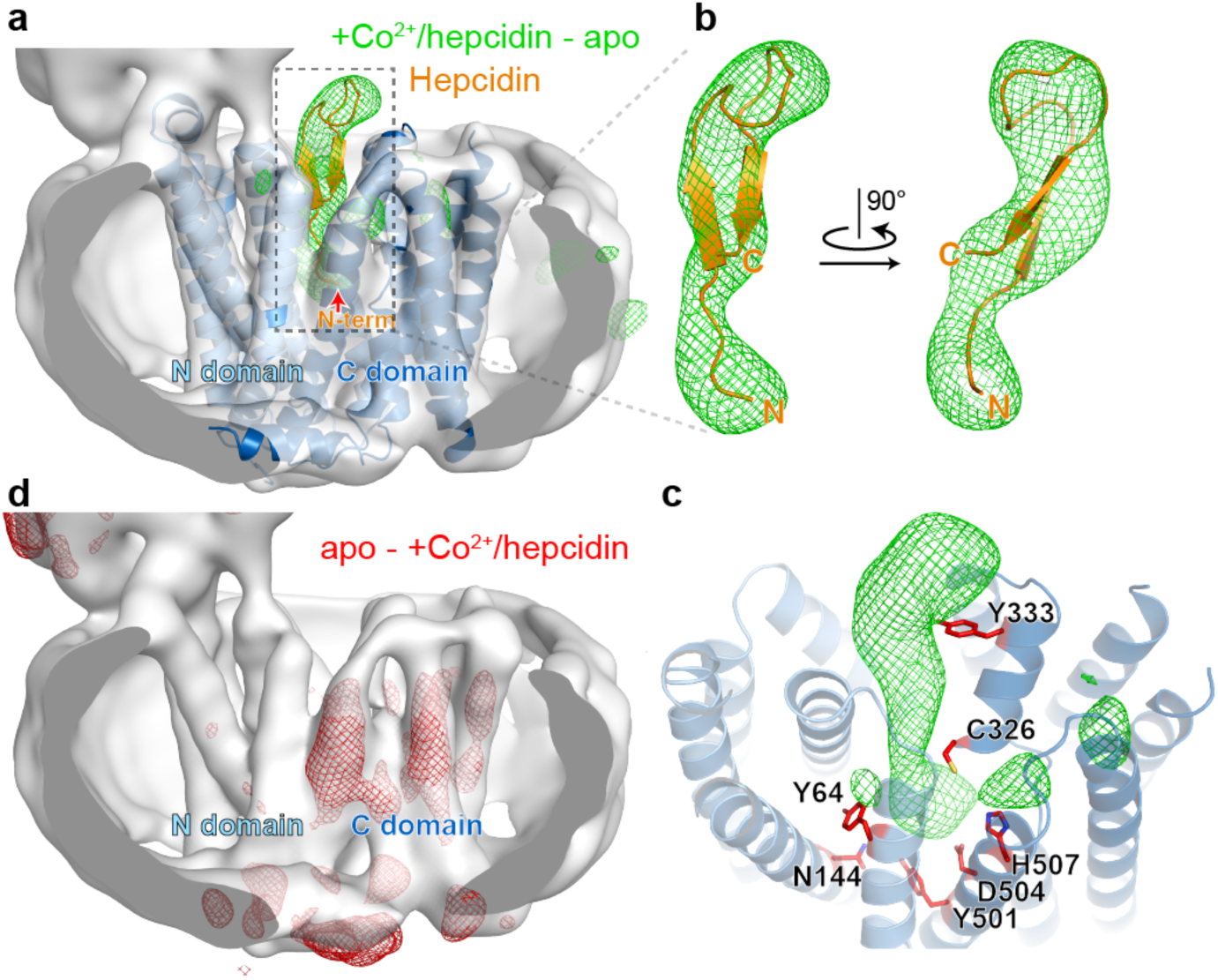
Hepcidin binding and FPN dynamics. **a**, Low pass filtered map for apo FPN (grey surface) with structural model of apo FPN (cartoon helices). Green mesh depicts difference density arising from subtracting the apo FPN map from that of FPN bound to Co^2+^ and hepcidin. Hepcidin from PDB ID: 3H0T was docked into the difference density. **b**, Close up views of speculative hepcidin pose within the difference density. The N-terminus is modeled as binding deep within the transmembrane cavity of FPN. **c**, Hepcidin-resistant FPN mutations leading to hemochromatosis (red sticks) map to regions surrounding the green difference density. **d**, Difference density after subtracting the Co^2+^ and hepcidin FPN map from the apo FPN map. The strongest difference peaks are with the C domain, indicating less ordered density for this region in the presence of Co^2+^ and hepcidin potentially arising from structural heterogeneity.

Despite our efforts to improve the resolution of the transmembrane domain of FPN:Co^2+^:hepcidin, the density of the FPN C domain is significantly weaker and less resolved than the N domain. The weaker C domain is also revealed in the difference map subtracting the FPN:Co^2+^:hepcidin map from the apo FPN map (Fig. 5d). The resolution difference between FPN N and C domains suggests increased conformational flexibility of the C domain in the presence of Co^2+^ and hepcidin. This disorder may itself lead to ubiquitination of FPN. Alternatively, conformational heterogeneity may produce a unique conformation required for ubiquitination.

## Discussion

Ferroportin is a central regulator of iron homeostasis in humans. Both human FPN and the bacterial homologue bbFPN show remarkable similarities in their overall architecture, with a unique conformation of TM7 likely responsible for molecular recognition of iron. This similarity suggests a deep evolutionary history for FPN-like transporters within the broader major facilitator superfamily. Hepcidin, by contrast, is specific to vertebrates, and likely evolved as a new strategy to regulate a critical point in iron absorption. Our structural analysis, combined with molecular dynamics simulations, and mutagenesis studies, map the location of iron and hepcidin binding to FPN.

We determined here that hepcidin binding to FPN is greatly potentiated by iron itself, potentially due to the stabilizing effect iron has on the hepcidin-binding site of FPN. With iron, the binding affinity of hepcidin falls in a range concordant with the concentration of hepcidin observed in healthy human adults. In normal iron homeostasis, this may enable hepcidin to selectively bind and regulate FPN molecules actively transporting iron and loaded with Fe^2+^, while sparing FPN molecules located on cells with low transport activity. Hepcidin binding to ferroportin would both trap the transporter in an outward open state and limit iron egress; both would acutely decrease iron efflux. Elevated hepcidin levels would inappropriately overcome this regulatory strategy and degrade FPN.

While the precise structural determinants underpinning hepcidin binding to FPN will require future studies capturing a high resolution hepcidin-ferroportin complex, the allosteric potentiation of hepcidin activity by iron may have immediate consequences for the development of hepcidin mimetics currently in clinical trials^16^. Furthermore, hepcidin antagonism by direct targeting of FPN may require molecules with high potency to overcome the nanomolar effect of the hormone in the presence of iron. The structural and functional insights into FPN function presented here therefore provide critical foundations for the discovery of therapeutics for human disorders of iron homeostasis.

## METHODS

No statistical methods were used to predetermine sample size. The experiments were not randomized and the investigators were not blinded to allocation during experiments and outcome assessment.

### Expression and purification of human ferroportin

The wild-type human FPN gene was cloned into a pVL1392 vector containing an expression cassette comprised of a C terminal human rhinovirus 3C (HRV-3C) protease recognition sequence followed by a human protein C epitope tag (EDQVDPRLIDGK) and an 8x polyhistidine tag. Baculovirus was generated using *Spodoptera frugiperda* Sf9 insect cells (unauthenticated and untested for mycoplasma contamination, Expression Systems 94-001F) and the construct was expressed in *Spodoptera frugiperda* Sf9 insect cells. Cells were collected 48 h after transduction and stored at −80°C until further use. Frozen cell pellets were thawed and washed with a hypotonic buffer (20 mM HEPES pH 7.50, 1 mM EDTA, supplemented with 20 µg/mL leupeptin, and 160 μg/mL benzamidine) before solubilizing with 50 mM HEPES pH 7.5, 300 mM NaCl, 1% (w/v) n-dodecyl-β-D-maltopyranoside (DDM, Anatrace), 0.1% (w/v) cholesteryl hemisuccinate (CHS, Steraloids), 1 mM EDTA supplemented with 20 µg/mL leupeptin, and 160 μg/mL benzamidine for 1 h at 4°C. Following centrifugation, the resulting supernatant was loaded on homemade anti-protein C antibody Sepharose beads and washed extensively in 50 mM HEPES pH 7.50, 300 mM NaCl, 2 mM CaCl_2_, 0.1% (w/v) DDM, 0.01% (w/v) CHS. FPN was eluted with 50 mM HEPES pH 7.50, 300 mM NaCl, 0.1% (w/v) DDM, 0.01% (w/v) CHS, 0.2 mg/mL Protein C peptide (Genscript) and 5 mM EDTA. The protein was concentrated with a Vivaspin 100-kDa MWCO concentrator and the monomeric FPN fraction was collected after size-exclusion chromatography (SEC) over a Superdex S200 Increase 10/300 GL column (GE Healthcare) equilibrated with 20 mM HEPES pH 7.50, 100 mM NaCl and 0.1% (w/v) DDM, and 0.01% (w/v) CHS.

### Expression and purification of MSP

Constructs encoding MSP-NW9 or MSP-NW11^39^ in the pET28b vector (Addgene #133442) were transformed into BL21(DE3) Rosetta *Escherichia coli*, and grown in terrific broth medium supplemented with 2 mM MgCl_2_ and 0.1% (w/v) glucose at 37°C. At OD_600_ of ∼0.6, expression was induced by addition of 400 µM isopropyl β-d-1-thiogalactopyranoside (IPTG) and lowering the temperature to 20°C. Cells were harvested after 16 hours and resuspended into 5 mL lysis buffer (200 mM Tris pH 8.0, 500 mM NaCl, 1% (v/v) Triton X-100 (Sigma), 0.02 mg/mL leupeptin, 0.16 mg/mL benzamidine, and benzonase) per gram pellet. After stirring for 30 min at 4°C, cells were lysed by pulsed sonication on ice. The lysate was cleared by centrifugation at 15,000 x g for 25 min at 4°C and loaded on Ni-NTA Sepharose. Ni-NTA beads were washed with 50 mM Tris pH 8.0, 500 mM NaCl, 1% (v/v) Triton, then 50 mM Tris pH 8.0, 500 mM NaCl, 50 mM sodium cholate, then 50 mM Tris pH 8.0, 500 mM NaCl, and finally with 50 mM Tris pH 8.0, 500 mM NaCl, 30 mM Imidazole. MSP was eluted with 50 mM Tris-HCl pH 8.0, 500 mM NaCl, 400 mM Imidazole and dialyzed into 50 mM Tris–HCl, pH 8.0, 100 mM NaCl, 1 mM EDTA, 0.1 mM TCEP at 4°C. The following day, MSP was concentrated on a Vivaspin 10-kDa MWCO concentrator, aliquots were flash frozen in liquid nitrogen and stored at -80°C for reconstitution.

### Isolation, expression and purification of Fab45D8

The heavy and light chain sequences of mAb45D8^25^ were separately cloned into pcDNA3.4 and the resulting vectors were transfected into Expi293F Human Embryonic Kidney cells (Life Technologies) using a 1:2 mass ratio of light and heavy chain DNA with the Expifectamine transfection kit (Life Technologies) as per the manufacturer’s instructions. Supernatant containing mAb45 was harvested 136 h after transfection and loaded on homemade Protein G Sepharose beads and extensively washed with a buffer comprising 20 mM HEPES pH 7.50, and 100 mM NaCl. mAb45D8 was eluted with 100 mM glycine (pH 3.0) and fractions were immediately neutralized with 200 mM HEPES pH 7.50. To generate the Fab fragment, 10 mg of purified mAb45D8.1 was diluted into 9.5 ml freshly prepared cleavage buffer (20 mM sodium phosphate pH 7.00, 10 mM EDTA, and 10 mM cysteine) and treated with 0.5 ml agarose immobilized papain (Thermo Scientific) at 37°C. After 16 h the cleaved Fab45D8 fragment was purified by reverse Protein A affinity chromatography, followed by SEC into buffer comprised of 20 mM HEPES pH 7.50 and 100 mM NaCl. Fab45D8 was concentrated on a Vivaspin 10-kDa MWCO concentrator, and aliquots were flash frozen in liquid nitrogen and stored at -80°C for later use.

### Reconstitution of FPN into lipidic nanodisc

Purified FPN (0.2-0.5 mg) was mixed with purified MSP and a lipid mixture containing a 2:3 weight ratio of 1-palmitoyl-2-oleoylphosphatidylcholine (POPC, Avanti) and 1-palmitoyl-2-oleoyl-sn-glycero-3-phospho-(1’-rac-glycerol) (POPG, Avanti). For reconstitution into NW9 nanodiscs, an FPN:MSP:Lipid molar ratio of 1:20:1100 was used. For reconstitution into NW11 nanodiscs, an FPN:MSP:Lipid molar ratio of 1:20:800 was used. The reconstitution sample was nutated for 1 h at 4°C before addition of 0.2 g/mL SM2-BioBeads (BioRad), and the reconstitution sample was further nutated overnight at 4°C before removal of the biobeads. FPN containing nanodiscs were purified by loading the reconstitution sample on anti-protein C antibody Sepharose beads and washing extensively with 20 mM HEPES pH 7.50, 100 mM NaCl, and 1 mM CaCl_2_ to remove empty nanodiscs. FPN containing nanodiscs were eluted with 20 mM HEPES pH 7.50, 100 mM NaCl, 0.25 mM EDTA, and 0.2 mg/mL Protein C peptide (Genscript), and concentrated on a Vivaspin 50-kDa concentrator.

### Crystallization and structure determination of Fab45D8

Purified Fab45D8 was diluted to 13.0 mg/mL in 20 mM HEPES pH 7.5, 100 mM NaCl. Fab45D8 crystals were obtained in 0.3 M trimethylamine-N-oxide (TMAO), 0.1 M Tris pH 8.5, and 30% (w/v) PEG 2000 MME at 20 °C. Individual crystals were flash frozen in liquid nitrogen after a 30 s soak in 0.3 M trimethylamine-N-oxide (TMAO), 0.1 M Tris pH 8.5, and 30% (w/v) PEG 2000, and 20% v/v ethylene glycol. A full diffraction dataset was collected at the Advanced Photon Source GM/CA-CAT beamline 23ID-B, and processed using xia2dials^40^ implementation of XDS^41^. The structure of the Fab was solved by molecular replacement using Phaser^42^, with a search model of a closely related germline mouse monoclonal antibody (PDB ID: 6BZV^43^) lacking complementarity determining regions (CDRs). The model was iteratively improved by refinement in Coot^44^ and Phenix^45^. Data collection and refinement statistics are summarized in Supplementary Table 1. The final model contained 96.77%, 2.23% and 0% in the favored, allowed and outlier regions of the Ramachandran plot, respectively as assessed by MolProbity^46^.

### Calcein transport assay for divalent cations

FPN was reconstituted into liposomes for divalent cation transport assays. Empty liposomes were prepared as a 3:1 mass ratio of 1-palmitoyl-2-oleoyl-sn-glycero-3-phosphoethanolamine (POPE, Avanti) to POPG dissolved in chloroform, followed by gentle evaporation of the chloroform under a stream of nitrogen gas, and overnight desiccation. The lipids were dissolved in 20 mM HEPES pH 7.40, 100 mM KCl to a final concentration of 12.5 mg/mL, sonicated until optically clear, subjected to multiple freeze-thaw cycles, and extruded through a 400 nm polycarbonate filter (Avestin) to generate unilamellar vesicles. Subsequently 0.13% (w/v) Triton-X100 (Sigma) was added to destabilize liposomes, corresponding to concentration yielding 80% of the maximum OD_540_ obtained in a liposome destabilization curve. Purified FPN was added at a 1:50 protein to lipid mass ratio and incubated for 15 min at 4°C. Control liposomes devoid of FPN were prepared in parallel using the same concentration of DDM. To remove excess detergent, 0.05 g/mL of SM2-BioBeads were added to the sample and nutated for 1 hr at 4°C, then 0.05 g/mL SM2-BioBeads were added followed by incubation overnight at 4°C, and finally addition of 0.08 g/mL SM2-BioBeads followed by incubation for 2 hr at 4°C. Proteoliposomes were harvested by ultracentrifugation at 300,000 x g for 30 min and resuspended at a concentration of 2.0 mg/mL lipids in internal buffer comprised of 20 mM HEPES pH 7.40 and 100 mM KCl, before flash freezing in liquid nitrogen and storage at -80°C. On the day of the transport assay, proteoliposomes were thawed and incubated with 500 mM calcein (Sigma), then subjected to three freeze-thaw cycles, and extruded through a 400 nm polycarbonate filter. The liposomes were washed four times with external buffer comprised of 20 mM HEPES pH 7.40 and 100 mM NaCl, by repeated ultracentrifugation and resuspension. Immediately prior to the assay, the calcein containing proteoliposomes were diluted to 0.25 mg/mL lipid in external buffer. Time-course fluorescence traces were recorded as 1 s integrations using a FluoroMax-4 (Horiba) with λ_ex_ of 490 nm and λ_em_ of 520 nm. Steady state fluorescence was recorded for at least 5 min, before addition of small aliquots of freshly prepared stocks of either FeCl_2_ or CoCl_2_. To stabilize the ferrous (Fe^2+^) state, we prepared iron as a 1:10 ratio of sodium ascorbate:FeCl2 immediately prior to the experiment. To determine the full extent of the calcein quenching response, 10 µM of the divalent cation ionophore calcimycin (Sigma) was added at the end of each experiment. Transport data was normalized to the mean baseline fluorescence intensity prior to addition of ion.

### Hepcidin binding assays

Fluorescence polarization measurements were performed using rhodamine-green labeled hepcidin (RhoG-hepcidin)^25^. For FPN saturation binding experiments, samples were prepared in a black 384-well plate (Greiner) containing 0-1 µM of nanodisc reconstituted NW11-FPN and 5 nM RhoG-hepcidin in sample buffer comprised of 20 mM HEPES pH 7.50, 100 mM NaCl, and supplemented with FeCl_2_, CoCl_2_ or MnCl_2_ as indicated. For ion stimulation experiments, 100 nM NW11-FPN and 5 nM RhoG-Hepcidin was mixed with 0-600 µM of CoCl_2_. For Fab binding experiments, 100 nM NW11-FPN and 5 nM RhoG-Hepcidin was mixed with 0 - 3 µM of Fab45D8 in sample buffer containing 10 µM CoCl_2._ Binding reactions were equilibrated for 60 min at RT, and fluorescence polarization was recorded on a Biotek Synergy H4 (Agilent) in polarization mode using fixed bandpass filters with λ_ex_ of 484 nm and λ_em_ of 520 nm.

Analytical fluorescence size exclusion chromatography (FSEC) was performed by mixing 25 µg of NW11-FPN with 2x fold molar excess of RhoG-Hepcidin in sample buffer comprised of 20 mM HEPES (pH 7.50), 100 mM NaCl and 10 µM CoCl_2_. Samples were incubated for 20 min on ice and 1.5 x molar excess of Fab45D8, or sample buffer, was added followed by incubation for 30 min on ice. Samples were injected on a Superdex 200 Increase 10/300 GL column (GE Lifesciences) pre-equilibrated in 20 mM HEPES pH 7.50, 100 mM NaCl, and 10 µM CoCl_2_. RhoG-Hepcidin fluorescence was recorded using an FP-1520 Intelligent Fluorescence Detector (Jasco) with λ_ex_ of 493 nm and λ_em_ of 524 nm.

### Cryo-EM Sample Preparation and Data Collection

Nanodisc-reconstituted apo FPN was mixed with 1.15 molar excess of Fab45D8 and incubated on ice for 30 min. The complex was purified by size-exclusion chromatography over a Superdex S200 Increase 10/300 GL column (GE Healthcare) equilibrated with 20 mM HEPES pH 7.50, 100 mM NaCl. For Co^2+^/hepcidin samples, 600 µM CoCl_2_ and 30 μM hepcidin (Bachem) was added to nanodisc-reconstituted FPN and incubated for 20 minutes on ice prior to addition of. Fab45D8. The resulting complex was purified over size-exclusion chromatography as for the apo sample but with the addition of 100 µM CoCl_2_ in the chromatography buffer. Collected fractions were supplemented with fresh hepcidin to 30 μM. For both preparations, fractions containing the nanodisc-FPN-Fab45D8 complex were concentrated to ∼3 mg/ml on a Vivaspin 50-kDa MWCO concentrator and freshly used for electron microscopy.

For high-resolution cryo-EM, the apo FPN-Fab45D8 complex was diluted to 0.0375 mg/mL in 20 mM HEPES pH 7.5, 100 mM NaCl directly prior to vitrification, and 2 μL sample was applied to glow-discharged gold holey carbon 1.2/1.3 300-mesh grids (Quantifoil) coated in-house with graphene oxide^47–49^. Grids were blotted for 2-4 seconds at 0 force and 10 seconds wait time before being plunge vitrified in liquid ethane using a MarkIV Vitrobot (ThermoFisher). The blotting chamber was maintained at 22°C and 100% humidity during freezing.

Co^2+^/hepcidin samples were diluted to 1.2 mg/mL in gel filtration buffer (20 mM HEPES pH 7.5, 100 mM NaCl, 100 µM CoCl_2_) before vitrification and 2 μL sample was applied to glow-discharged UltrAufoil 1.2/1.3 300-mesh grids (Quantifoil). Grids were blotted for 3 seconds at 0 force and 5 seconds wait time before being plunge vitrified in liquid ethane using a MarkIV Vitrobot (ThermoFisher). The blotting chamber was maintained at 22°C and 100% humidity during freezing.

FPN-Fab45D8 movies were collected using a Titan Krios (ThermoFisher) outfitted with a K3 camera and Bioquantum energy filter (Gatan). The K3 detector was operated in superresolution mode and the energy filter slit width was set to 20 eV. Movies were collected at a nominal magnification of 105,000x, physical pixel size 0.834Å/pix, with a 70 µm C2 aperture and 100 µm objective aperture at a dose rate of 8 e^-^/pixel/second. A total dose of 66 e^-^/Å^2^ was collected as a 120-frame movie, resulting in a 6-second movie with 0.55 e^-^/frame. All data were collected using semi-automated imaging scripts in SerialEM^50^. 5009 movies were collected using a 3×3 image shift pattern at 0° tilt and 406 movies were collected on-axis with a 30° stage tilt in two separate data collection sessions.

Co2+/hepcidin-FPN-Fab45D8 movies were collected using a Talos Arctica (ThermoFisher) outfitted with a K3 camera (Gatan). The K3 detector was operated in superresolution mode and movies were collected at a nominal magnification of 28,000x, superresolution pixel size 0.7 Å/pix, with a 50 µm C2 aperture and 100 µm objective aperture at a dose rate of 10 e^-^/pixel/second. A total dose of 60 e^-^/Å^2^ was collected as a 120-frame movie, resulting in a 11.5-second movie with 0.5 e^-^/frame. All data were collected using semi-automated imaging scripts in SerialEM^50^. 958 movies were collected using a 3×3 image shift pattern at 0° tilt.

### Cryo-EM Image Processing

For FPN-Fab45D8, data were motion corrected and 2x binned on-the-fly using MotionCor2^51^ in the SCIPION pipeline^52^. Motion corrected micrographs were imported into cryoSPARC^53^ and RELION^54^ and contrast transfer function parameters were calculated using CTFFIND4^55^. CTF information for tilted images were estimated using patch CTF estimation in cryoSPARC. 138,314 particles were selected from 497 micrographs using the Blob picker in cryoSPARC. 2D class averages were generated after extracting the putative particles with a 300-pixel box and binning to 64 pixels. Six of these averages were used as templates for further particle picking. Template picking yielded 4,737,795 particles. These were split into 6 groups to increase speed of processing, extracted in a 300-pixel box, and binned to 64 pixels. 2D classification was run in cryoSPARC with default settings except: number of 2D classes 200, initial classification uncertainty factor 4, number of online-EM iterations 40, batch size per class 300. Objectively “good” (showing clear Fab and receptor density) class averages were selected and exported to RELION format using csparc.py^56^. Class averages that were not classified as “good”, but were not clearly ice contamination or graphene oxide edges, were run through a second round of 2D classification with default settings except: number of 2D classes 200, number of online-EM iterations 40, batch size per class 300. All “saved” class averages from the second rounds of 2D classification in cryoSPARC selected and exported to RELION format using csparc.py. Particles were extracted from CTF-corrected images in RELION at a box size of 300 pixels, binned to 128 pixels. 1,326,130 particles, in three groups (detailed in Supplementary Fig. 3) were classified in 3D with image alignment in RELION using an initial model generated in cryoSPARC from 80,000 particles collected on a Talos Arctica filtered to 40 Å, C1 symmetry, a regularization parameter of 4, for 30-35 iterations with no mask. Particles from classes with resolved transmembrane (TM) helices were selected, extracted in a 300-pixel box, and imported back into cryoSPARC. Non-uniform refinement was run with default settings and no resolution limit, resulting in angle and shift assignments for 850,000 particles. These particles were subsequently exported to RELION format using csparc.py and run through 3D classification without image alignment in RELION. Four of the 12 classes were selected, imported into RELION, run through non-uniform alignment with an automatically generated mask, and refined to a reported global resolution of 3.2 Å. The resulting map showed clear signs of mild preferred orientation (Supplementary Fig. 3). The particles were exported into RELION format using csparc.py, converted into an image stack, and imported into cisTEM^57^ as a refinement package. The particles were reconstructed and half-maps were generated using the “generate 3D” command. These half maps, as well as the half maps from cryoSPARC were run through our lab’s directional Fourier shell correlation (dFSC) program^58^ clearly showing a more distributed range of views in the map generated by cisTEM. Maps were sharpened in RELION. Resolutions are reported using the FSC = 0.143 cut-off^59^ and were estimated in cryoSPARC and cisTEM.

For Co2+/hepcidin-FPN-Fab45D8, data were motion corrected without binning on-the-fly using MotionCor2^51^ in the SCIPION pipeline^52^. Motion corrected micrographs were imported into cryoSPARC^53^ and contrast transfer function parameters were calculated using CTFFIND4^55^. 562,839 particles were selected from 598 micrographs using the Blob picker in cryoSPARC. 2D class averages were generated after extracting the putative particles with a 360-pixel box and binning to 64 pixels. Nine of these averages were used as templates for further particle picking. Template picking yielded 1,356,071 particles. These particles were extracted in a 360-pixel box, and binned to 64 pixels. 2D classification was run in cryoSPARC with default settings except: number of 2D classes 200, initial classification uncertainty factor 4, number of online-EM iterations 40, batch size per class 300. Objectively “good” (showing clear Fab and receptor density) class averages were selected for 3D classification. Class averages that were not classified as “good”, but were not clearly ice contamination or graphene oxide edges, were run through a second round of 2D classification with default settings except: number of 2D classes 200, number of online-EM iterations 40, batch size per class 300. All “saved” class averages from the second rounds of 2D classification in cryoSPARC were sent to 3D classification. These particles were subjected to two rounds of heterogeneous refinement in cryoSPARC that serves as a “trash collector”. Four initial models were used, three generated from an early round of ab initio model generation and our final apo FPN-Fab45D8 structure. All initial models were filtered to 30Å before refinement. Particles were unbinned and a final heterogeneous refinement was performed with three good initial models of apo FPN-Fab45D8. Non-uniform refinement was run with default settings and no resolution limit, on the class showing the clearest density in the outward-open cavity next to helix 7b resulting in angle and shift assignments for 57,669 particles. Our in-house directional fourier shell correlation (dFSC) program^58^ was run on the halfmaps generated from this refinement. Resolutions are reported using the FSC = 0.143 cut-off^59^ and were estimated in cryoSPARC.

### Difference Map Generation

A density map was generated in Chimera^60^ (molmap) around the receptor of our FPN-Fab45D8 model. Density was erased for all parts of the protein except for a color zone area 2 Å wide around helices H1, H3, H4, and H6. This density was resampled onto the grid of our highest resolution map (EMD-21539) and saved as a new map of the “N-terminal helices”. Our apo and hepcidin bound structures (EMD-21539 and EMD-21550) were aligned to the N-terminal helices using “Fit to Map” in Chimera with the criteria of correlation selected. These aligned maps were then resampled onto the same grid as the N-terminal helices. The two aligned and resampled maps were filtered to 10 Å using e2proc3d.py^61^. These maps were then used to generate difference maps (apo-hep and hep-apo) using diffmap.exe^62^. These difference maps were visualized in Chimera along with the 10 Å filtered apo map for comparison.

### Model building and refinement

A homology model of human FPN in the outward open state was built using Modeller^63^, with a previously determined X-ray crystal structure of outward open bbFPN (PDB ID: 5AYN)^23^ as a template. After truncating putatively flexible regions (N-terminus, ICL3, ECL5, and C-terminus), the resulting model was fit into the 3.2 Å cryo-EM map of FPN:Fab45D8 using Chimera^60^. The initial template was manually rebuilt in Coot^44^ and iteratively refined with real space refinement implemented in Phenix^45^. Model geometry was assessed using MolProbity^46^. Further validation was performed with EMRinger^64^ to compare the map and final model. Map-to-model FSCs were calculated within Phenix. Figures were prepared in Chimera^60^ and PyMol.

To dock hepcidin into the Co^2+^/hepcidin-bound and apo FPN difference density, we used a previously determined X-ray crystal structure of hepcidin bound to a neutralizing Fab as a starting model^21^. Hepcidin, without the first two residues, was manually placed within the difference density in Coot, then real-space refined to conform to the difference density with maintaining the disulfide connectivity and secondary structure observed in the starting model. The resulting model docked to FPN has an overall RMSD of 1.2 Å compared to the starting model for regions with defined secondary structure.

### Molecular dynamics simulations

The structure of the outward-open apo conformation of human FPN was used as the starting coordinates for all simulations. Three different conditions were simulated (Supplementary Table 3): (1) the iron-absent condition, where no iron was added; (2) the iron-bound condition, where an Fe^2+^ ion was placed in the proposed iron binding site 5.6 Å from D325 α-carbon, 6.9 Å from D504 α-carbon, and 7.8 Å from H507 α-carbon; (3) the iron-in-bulk-solvent condition, where 15 Fe^2+^ ions were placed randomly in the water box outside the protein using Dabble^65^.

Simulation coordinates were prepared by removing non-FPN molecules from the initial structure. Prime (Schrödinger) was used to model missing side chains, and neutral acetyl and methylamide groups were added to cap protein termini. The unresolved loops between TM6-TM7 and TM9-TM10 (residues 239-290 and 394-451 respectively, inclusively) were not modeled. The termini surrounding these loops were capped. PropKa was used to determine the dominant protonation state of all titratable residues at pH 7^66,67^. The structure was internally hydrated using Dowser^68^. Dabble was used to additionally fill the extracellular cavity^65^. The structure was aligned using the Orientation of Proteins in Membranes (OPM) server^69^.

Using Dabble, the protein was inserted into a pre-equilibrated 1-palmitoyl-2-oleoylphosphatidylcholine (POPC) membrane bilayer. For all simulations except condition three (iron in bulk solvent), sodium and chloride ions were added at 150 mM to neutralize the system. For condition three, chloride ions were added to neutralize the system resulting in a concentration of 108 mM. A periodic box was used with dimensions 90 x 90 Å in the x-y plane and a water buffer of 10 Å above and below the protein to the periodic boundary. We used the CHARMM36m parameters for lipids, proteins, sodium and chloride ions, and the TIP3P model for waters^70–72^. The Fe^2+^ Lennard-Jones parameters were obtained from Li *et. al.’s* compromise model^73^.

All simulations were run on a single Graphical Processing Unit (GPU) using the Amber18 Compute Unified Device Architecture (CUDA) version of particle-mesh Ewald molecular dynamics (PMEMD)^74,75^. For each condition, 6 replicates were run. For each independent replicate, the system was minimized with 500 steps of steepest descent followed by 500 steps of conjugate gradient descent three times. 10 and 5 kcal mol^-1 -^Å^2^ harmonic restraints were used on the protein, lipid, and Fe^2+^ ions for the first and second minimization, respectively. 1 kcal mol^-1 -^Å^2^ harmonic restraints were used on the protein and Fe^2+^ ions for the final minimization. The system was then heated from 0 K to 100 K over 12.5 ps in the NVP ensemble with a Langevin thermostat and harmonically restraining the protein heavy atoms and Fe^2+^ ions with a restraint of 10 kcal mol^-1 -^Ȧ^2^. The system was further heated with the same restraints from 100 K to 310 K in the NPT ensemble over 125 ps. The system was equilibrated with harmonic restraints on protein heavy atoms and Fe^2+^ ions for 30 ns. The restraint strength started at 5 kcal mol^-1 -^Ȧ^2^ and was reduced by 1 kcal mol^-1 -^Å^2^ every 2 ns for the first 10 ns and then by 0.1 kcal mol^-1 -^Ȧ^2^ every 2 ns for the final 20 ns. Production simulations were performed at 310 K and 1 bar using the NPT ensemble, a Langevin thermostat and a Monte Carlo barostat. Every 200 ps snapshots were saved. All simulations were run for at least 2.2 μs. These simulations used a 4-fs time step with hydrogen mass repartitioning^76^. Bond lengths to hydrogen atoms were constrained using SHAKE^76,77^. Non-bonded interactions were cut off at 9 Å.

### Simulation Analysis Methods

MD snapshots were reimaged every 1 ns and centered using CPPTRAJ package in AmberTools18^78^. Simulations were visualized using Visual Molecular Dynamics and figures prepared in PyMOL^79^. Time traces from simulation were smoothed using a moving average with a window size of 15 ns unless otherwise indicated and visualized with the PyPlot package from Matplotlib. For all analysis in the manuscript that required structural alignment, we aligned to the initial Ferroportin structure using the backbone atoms of residues 26-116, 127-228, 308-483, and 492-543.

To investigate the localization of Fe^2+^ ions, the iron-in-bulk-solvent simulations (condition 3) were analyzed. To visualize the density of Fe^2+^ ions, the position of Fe^2+^ ions was recorded every 10 ns for each of the 6 simulation replicates, each 2 μs in length. Each Fe^2+^ ion position was then drawn as a point superimposed on the starting structure (Fig. 2b). To quantify the binding events, the distance between iron and the closest side chain oxygen atom on D325 was measured. This distance was graphed over 1.2 μs, including the equilibration time (Fig. 2c). A frame at 760 ns of an iron-bound simulation (condition 2) was selected to display an observed coordination state of the iron in the binding pocket (Fig. 2d). Note that other coordination arrangements were also observed. The persistent Fe^2+^ interactions are quantified in Supplementary Fig. 6. For each condition, the first 2 μs of each simulation, excluding equilibration, was selected. The distances measured were Fe^2+^-H507:NE2, Fe^2+^-D325:OD1/OD2, Fe^2+^-D504:OD1/OD2, and Fe^2+^-T320:O. The direct and near interaction frequencies were calculated as the fraction of the 2 μs that a Fe^2+^ ion was within 3 Å or 5.5 Å of the specified atoms. To assess the fraction of time the iron ions were directly interacting with or near the proposed binding site, the portion of time for which any of the previously mentioned interactions were direct or near was calculated.

To investigate the dynamics and conformation of the TM7b region, the iron-absent (condition 1) and iron-bound (condition 2) simulations were compared. The TM1-TM7b distance was measured using distance between the Cα of V51 and the Cα of Y333. For each simulation, we calculated the average of the distance over 2.2 μs, excluding equilibration. The average over the simulations for each condition was plotted with error bars representing the standard error of the mean (s.e.m.) (Fig. 4e right). The dynamics of D325 were also investigated. The dynamics were visualized by overlaying representative frames showing the movement of D325, the binding site, and TM7b. For iron-absent simulations, frames from a single replicate at 200, 350, 500, and 550 ns were overlaid (Fig. 4b). For iron-bound simulations, frames from a single replicate at 200, 500, 725, 1000 ns were overlaid (Fig. 4c). The conformational range of D325 was quantified by measuring the distance between Cγ of D325 and Cβ of S47. This was visualized for one replicate for each condition over a time of 1 μs inclusive of equilibration (Fig. 4d). For each independent replicate, the mean of the Cγ D325 - Cβ S47 distance was calculated over 2.2 μs. For each condition, the average over the replicates was plotted with error bars representing the s.e.m. (Fig. 4e left). The flexibility of D325 was quantified by calculating the root-mean-square fluctuation (RMSF) of the side-chain atoms of D325 using an in-house script (Fig 4e. center). Statistical significance was determined using the Mann-Whitney U test.

## Data Availability

All data generated or analyzed during this study are included in this published article and its Supplementary Information. Crystallographic coordinates and structure factors for the Fab45D8 complex have been deposited in the Protein Data Bank under accession code 6W4V. Coordinates for Fab45D8-FPN complex have been deposited in the Protein Data Bank under accession code 6W4S and the maps have been deposited in the Electron Microscopy Data Bank under accession code 21539. Maps for the FPN-Co^2+^-hepcidin-Fab45D8 complex have been deposited in the Electron Microscopy Data Bank under accession code 21550.

## REFERENCES

1. Drakesmith, H., Nemeth, E. & Ganz, T. Ironing out Ferroportin. Cell Metab. 22, 777–787 (2015).

2. Donovan, A. et al. The iron exporter ferroportin/Slc40a1 is essential for iron homeostasis. Cell Metab. 1, 191–200 (2005).

3. Donovan, A. et al. Positional cloning of zebrafish ferroportin1 identifies a conserved vertebrate iron exporter. Nature 403, 776–781 (2000).

4. Knutson, M. D., Oukka, M., Koss, L. M., Aydemir, F. & Wessling-Resnick, M. Iron release from macrophages after erythrophagocytosis is up-regulated by ferroportin 1 overexpression and down-regulated by hepcidin. Proc. Natl. Acad. Sci. U. S. A. 102, 1324–1328 (2005).

5. Ward, D. M. & Kaplan, J. Ferroportin-mediated iron transport: expression and regulation. Biochim. Biophys. Acta 1823, 1426–1433 (2012).

6. Aschemeyer, S. et al. Structure-function analysis of ferroportin defines the binding site and an alternative mechanism of action of hepcidin. Blood, The Journal of the American Society of Hematology 131, 899–910 (2018).

7. Nemeth, E. et al. Hepcidin regulates cellular iron efflux by binding to ferroportin and inducing its internalization. Science 306, 2090–2093 (2004).

8. De Domenico, I. et al. The molecular mechanism of hepcidin-mediated ferroportin down-regulation. Mol. Biol. Cell 18, 2569–2578 (2007).

9. Qiao, B. et al. Hepcidin-induced endocytosis of ferroportin is dependent on ferroportin ubiquitination. Cell Metab. 15, 918–924 (2012).

10. Ross, S. L. et al. Molecular mechanism of hepcidin-mediated ferroportin internalization requires ferroportin lysines, not tyrosines or JAK-STAT. Cell Metab. 15, 905–917 (2012).

11. Roetto, A. et al. Mutant antimicrobial peptide hepcidin is associated with severe juvenile hemochromatosis. Nat. Genet. 33, 21–22 (2003).

12. De Domenico, I. et al. The molecular basis of ferroportin-linked hemochromatosis. Proc. Natl. Acad. Sci. U. S. A. 102, 8955–8960 (2005).

13. Drakesmith, H. et al. Resistance to hepcidin is conferred by hemochromatosis-associated mutations of ferroportin. Blood 106, 1092–1097 (2005).

14. Roy, C. N. et al. Hepcidin antimicrobial peptide transgenic mice exhibit features of the anemia of inflammation. Blood 109, 4038–4044 (2007).

15. Ganz, T. & Nemeth, E. The hepcidin-ferroportin system as a therapeutic target in anemias and iron overload disorders. Hematology Am. Soc. Hematol. Educ. Program 2011, 538–542 (2011).

16. Manolova, V. et al. Oral ferroportin inhibitor ameliorates ineffective erythropoiesis in a model of β-thalassemia. J. Clin. Invest. (2019) doi: 10.1172/JCI129382.

17. Witcher, D. R. et al. LY2928057, an antibody targeting ferroportin, is a potent inhibitor of hepcidin activity and increases iron mobilization in normal cynomolgus monkeys. (2013).

18. Crielaard, B. J., Lammers, T. & Rivella, S. Targeting iron metabolism in drug discovery and delivery. Nat. Rev. Drug Discov. 16, 400–423 (2017).

19. Vlasveld, L. T. et al. Twenty Years of Ferroportin Disease: A Review or An Update of Published Clinical, Biochemical, Molecular, and Functional Features. Pharmaceuticals 12, (2019).

20. Nemeth, E. et al. The N-terminus of hepcidin is essential for its interaction with ferroportin: structure-function study. Blood 107, 328–333 (2006).

21. Jordan, J. B. et al. Hepcidin revisited, disulfide connectivity, dynamics, and structure. J. Biol. Chem. 284, 24155–24167 (2009).

22. Bonaccorsi di Patti, M. C. et al. A structural model of human ferroportin and of its iron binding site. FEBS J. 281, 2851–2860 (2014).

23. Taniguchi, R. et al. Outward- and inward-facing structures of a putative bacterial transition-metal transporter with homology to ferroportin. Nat. Commun. 6, 8545 (2015).

24. Deshpande, C. N. et al. Calcium is an essential cofactor for metal efflux by the ferroportin transporter family. Nat. Commun. 9, 3075 (2018).

25. Ross, S. L. et al. Identification of Antibody and Small Molecule Antagonists of Ferroportin-Hepcidin Interaction. Front. Pharmacol. 8, 838 (2017).

26. Wu, S. et al. Fabs enable single particle cryoEM studies of small proteins. Structure 20, 582–592 (2012).

27. Guellec, J. et al. Molecular model of the ferroportin intracellular gate and implications for the human iron transport cycle and hemochromatosis type 4A. FASEB J. 33, 14625–14635 (2019).

28. Wang, J. On the appearance of carboxylates in electrostatic potential maps. Protein Sci. 26, 396–402 (2017).

29. White, M. D. & Flashman, E. Catalytic strategies of the non-heme iron dependent oxygenases and their roles in plant biology. Curr. Opin. Chem. Biol. 31, 126–135 (2016).

30. Sham, R. L. et al. Autosomal dominant hereditary hemochromatosis associated with a novel ferroportin mutation and unique clinical features. Blood Cells Mol. Dis. 34, 157–161 (2005).

31. Fernandes, A. et al. The molecular basis of hepcidin-resistant hereditary hemochromatosis. Blood, The Journal of the American Society of Hematology 114, 437–443 (2009).

32. Galesloot, T. E. et al. Serum hepcidin: reference ranges and biochemical correlates in the general population. Blood 117, e218–25 (2011).

33. Mitchell, C. J., Shawki, A., Ganz, T., Nemeth, E. & Mackenzie, B. Functional properties of human ferroportin, a cellular iron exporter reactive also with cobalt and zinc. Am. J. Physiol. Cell Physiol. 306, C450–9 (2014).

34. Madejczyk, M. S. & Ballatori, N. The iron transporter ferroportin can also function as a manganese exporter. Biochim. Biophys. Acta 1818, 651–657 (2012).

35. Yin, Z. et al. Ferroportin is a manganese-responsive protein that decreases manganese cytotoxicity and accumulation. J. Neurochem. 112, 1190–1198 (2010).

36. Jin, L. et al. Mice overexpressing hepcidin suggest ferroportin does not play a major role in Mn homeostasis. Metallomics 11, 959–967 (2019).

37. Clark, R. J. et al. Understanding the structure/activity relationships of the iron regulatory peptide hepcidin. Chem. Biol. 18, 336–343 (2011).

38. Preza, G. C. et al. Minihepcidins are rationally designed small peptides that mimic hepcidin activity in mice and may be useful for the treatment of iron overload. J. Clin. Invest. 121, 4880–4888 (2011).

39. Nasr, M. L. et al. Covalently circularized nanodiscs for studying membrane proteins and viral entry. Nat. Methods 14, 49–52 (2017).

40. Winter, G., Lobley, C. M. C. & Prince, S. M. Decision making in xia2. Acta Crystallogr. D Biol. Crystallogr. 69, 1260–1273 (2013).

41. Kabsch, W. XDS. Acta Crystallogr. D Biol. Crystallogr. 66, 125–132 (2010).

42. McCoy, A. J. et al. Phaser crystallographic software. J. Appl. Crystallogr. 40, 658–674 (2007).

43. Aleman, F. et al. Immunogenetic and structural analysis of a class of HCV broadly neutralizing antibodies and their precursors. Proc. Natl. Acad. Sci. U. S. A. 115, 7569–7574 (2018).

44. Emsley, P. & Cowtan, K. Coot: model-building tools for molecular graphics. Acta Crystallogr. D Biol. Crystallogr. 60, 2126–2132 (2004).

45. Adams, P. D. et al. PHENIX: a comprehensive Python-based system for macromolecular structure solution. Acta Crystallogr. D Biol. Crystallogr. 66, 213–221 (2010).

46. Chen, V. B. et al. MolProbity: all-atom structure validation for macromolecular crystallography. Acta Crystallogr. D Biol. Crystallogr. 66, 12–21 (2010).

47. Cote, L. J., Kim, F. & Huang, J. Langmuir-Blodgett assembly of graphite oxide single layers. J. Am. Chem. Soc. 131, 1043–1049 (2009).

48. Palovcak, E. et al. A simple and robust procedure for preparing graphene-oxide cryo-EM grids. J. Struct. Biol. 204, 80–84 (2018).

49. Wang, F. et al. Amino and PEG-amino graphene oxide grids enrich and protect samples for high-resolution single particle cryo-electron microscopy. J. Struct. Biol. 209, 107437 (2020).

50. Mastronarde, D. N. SerialEM: A Program for Automated Tilt Series Acquisition on Tecnai Microscopes Using Prediction of Specimen Position. Microsc. Microanal. 9, 1182–1183 (2003).

51. Zheng, S. Q. et al. MotionCor2: anisotropic correction of beam-induced motion for improved cryo-electron microscopy. Nat. Methods 14, 331–332 (2017).

52. de la Rosa-Trevín, J. M. et al. Scipion: A software framework toward integration, reproducibility and validation in 3D electron microscopy. J. Struct. Biol. 195, 93–99 (2016).

53. Punjani, A., Rubinstein, J. L., Fleet, D. J. & Brubaker, M. A. cryoSPARC: algorithms for rapid unsupervised cryo-EM structure determination. Nat. Methods 14, 290–296 (2017).

54. Scheres, S. H. W. RELION: implementation of a Bayesian approach to cryo-EM structure determination. J. Struct. Biol. 180, 519–530 (2012).

55. Rohou, A. & Grigorieff, N. CTFFIND4: Fast and accurate defocus estimation from electron micrographs. J. Struct. Biol. 192, 216–221 (2015).

56. Asarnow, D., Palovcak, E. & Cheng, Y. asarnow/pyem: UCSF pyem v0.5. (2019). doi: 10.5281/zenodo.3576630.

57. Grant, T., Rohou, A. & Grigorieff, N. cisTEM, user-friendly software for single-particle image processing. Elife 7, (2018).

58. Dang, S. et al. Cryo-EM structures of the TMEM16A calcium-activated chloride channel. Nature 552, 426–429 (2017).

59. Rosenthal, P. B. & Henderson, R. Optimal determination of particle orientation, absolute hand, and contrast loss in single-particle electron cryomicroscopy. J. Mol. Biol. 333, 721–745 (2003).

60. Goddard, T. D., Huang, C. C. & Ferrin, T. E. Visualizing density maps with UCSF Chimera. J. Struct. Biol. 157, 281–287 (2007).

61. Tang, G. et al. EMAN2: an extensible image processing suite for electron microscopy. J. Struct. Biol. 157, 38–46 (2007).

62. diffmap | The Grigorieff Lab. https://grigoriefflab.umassmed.edu/diffmap.

63. Eswar, N. et al. Comparative protein structure modeling using MODELLER. Curr. Protoc. Protein Sci. Chapter 2, Unit 2.9 (2007).

64. Barad, B. A. et al. EMRinger: side chain-directed model and map validation for 3D cryo-electron microscopy. Nat. Methods 12, 943–946 (2015).

65. Betz, R. Dabble. (2017). doi: 10.5281/zenodo.836914.

66. Olsson, M. H. M., Søndergaard, C. R., Rostkowski, M. & Jensen, J. H. PROPKA3: Consistent Treatment of Internal and Surface Residues in Empirical pKa Predictions. J. Chem. Theory Comput. 7, 525–537 (2011).

67. Søndergaard, C. R., Olsson, M. H. M., Rostkowski, M. & Jensen, J. H. Improved Treatment of Ligands and Coupling Effects in Empirical Calculation and Rationalization of pKa Values. J. Chem. 66. Theory Comput. 7, 2284–2295 (2011).

68. Morozenko, A. & Stuchebrukhov, A. A. Dowser, a new method of hydrating protein structures. Proteins: Structure, Function, and Bioinformatics vol. 84 1347–1357 (2016).

69. Lomize, M. A., Lomize, A. L., Pogozheva, I. D. & Mosberg, H. I. OPM: orientations of proteins in membranes database. Bioinformatics 22, 623–625 (2006).

70. Klauda, J. B. et al. Update of the CHARMM all-atom additive force field for lipids: validation on six lipid types. J. Phys. Chem. B 114, 7830–7843 (2010).

71. Huang, J. et al. CHARMM36m: an improved force field for folded and intrinsically disordered proteins. Nat. Methods 14, 71–73 (2017).

72. Beglov, D. & Roux, B. Finite representation of an infinite bulk system: Solvent boundary potential for computer simulations. J. Chem. Phys. 100, 9050–9063 (1994).

73. Li, P., Roberts, B. P., Chakravorty, D. K. & Merz, K. M., Jr. Rational design of particle mesh Ewald compatible Lennard-Jones parameters for+ 2 metal cations in explicit solvent. J. Chem. Theory Comput. 9, 2733–2748 (2013).

74. Case, D. A. et al. AMBER 2018; 2018. University of California, San Francisco.

75. Salomon-Ferrer, R., Götz, A. W., Poole, D., Le Grand, S. & Walker, R. C. Routine Microsecond Molecular Dynamics Simulations with AMBER on GPUs. 2. Explicit Solvent Particle Mesh Ewald. J. Chem. Theory Comput. 9, 3878–3888 (2013).

76. Hopkins, C. W., Le Grand, S., Walker, R. C. & Roitberg, A. E. Long-Time-Step Molecular Dynamics through Hydrogen Mass Repartitioning. J. Chem. Theory Comput. 11, 1864–1874 (2015).

77. Ryckaert, J.-P., Ciccotti, G. & Berendsen, H. J. C. Numerical integration of the cartesian equations of motion of a system with constraints: molecular dynamics of n-alkanes. J. Comput. Phys. 23, 327–341 (1977).

78. Roe, D. R. & Cheatham, T. E., 3rd. PTRAJ and CPPTRAJ: Software for Processing and Analysis of Molecular Dynamics Trajectory Data. J. Chem. Theory Comput. 9, 3084–3095 (2013).

79. Humphrey, W., Dalke, A. & Schulten, K. VMD: visual molecular dynamics. J. Mol. Graph. 14, 33–8, 66. 27–8 (1996).

