## Supplementary Information for "Structure of human ferroportin reveals molecular basis of iron homeostasis"

### SUPPLEMENTARY FIGURES

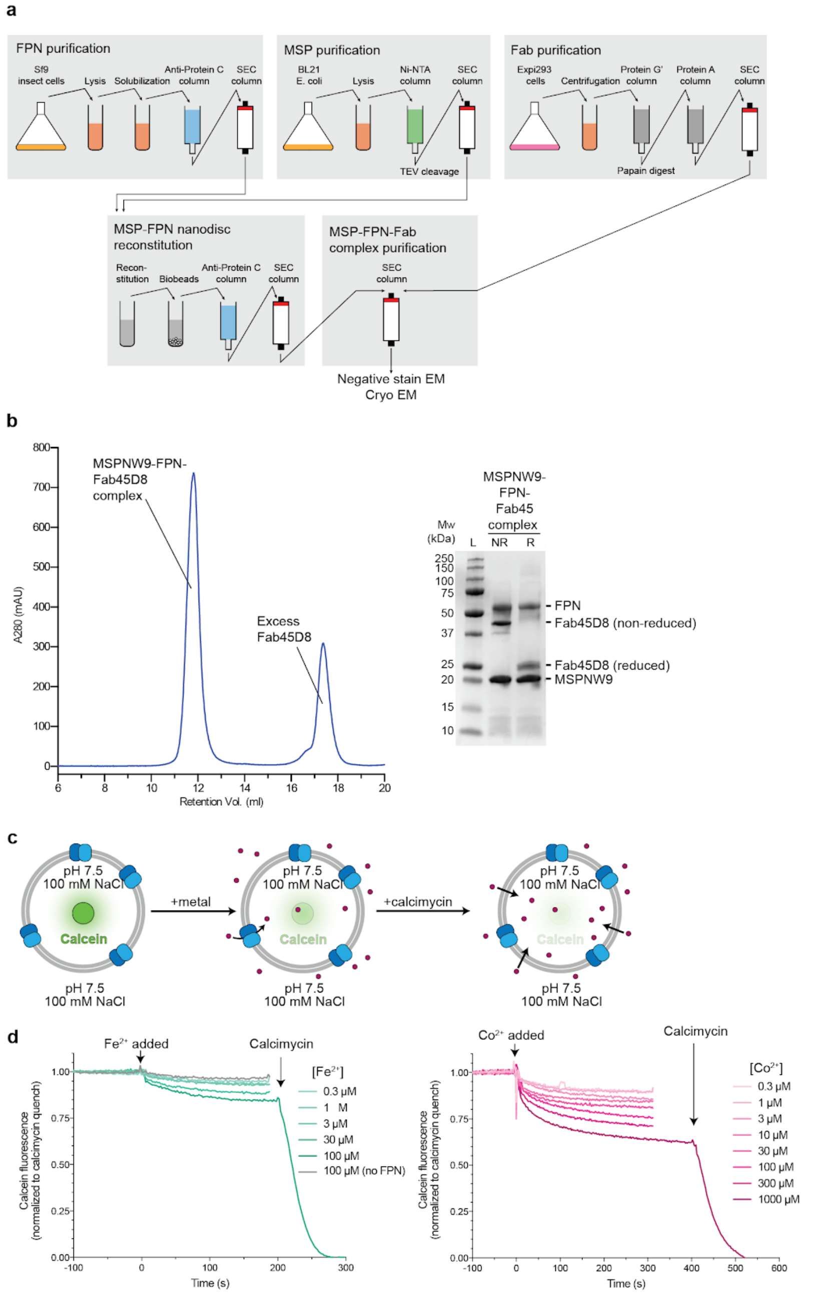


**Supplementary Figure 1. Biochemistry of purified human ferroportin. a,** Purification scheme for nanodisc-reconstituted FPN bound to Fab45D8 for cryo-EM studies. b, Size exclusion chromatography of purified FPN-Fab45D8 complex. SDS-PAGE gel of purified complex under non-reducing (NR) and reducing (R) conditions. **c,** Schematic of calcein-based assay to measure transport of divalent cations. Purified FPN is reconstituted into proteoliposomes containing calcein. Addition of divalent cations leads to quenching of calcein fluorescence. Addition of the divalent cation ionophore calcimycin fully quenches calcein fluorescence. **d,** Reconstituted human FPN transports $Fe^{2+}$ and $Co^{2+}$.

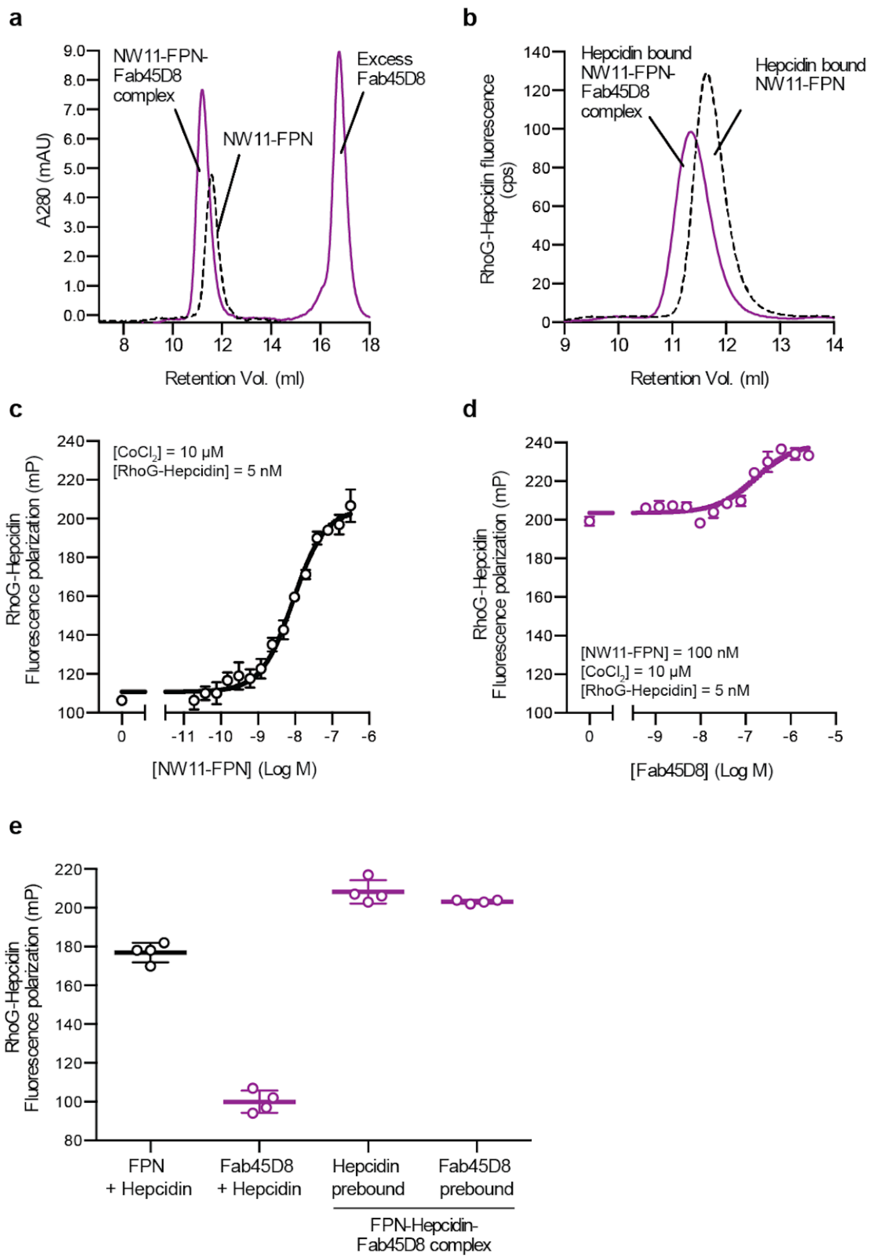


**Supplementary Figure 2. Characterization of Fab45D8. a,** Fab45D8 binds nanodisc-reconstituted FPN (NW11-FPN), as assessed by size-exclusion chromatography. **b,** Fluorescence size exclusion chromatography of rhodamine G hepcidin (RhoG-hepcidin). RhoG-hepcidin co-elutes with NW11-FPN and with a NW11-FPN:Fab45D8 complex. **c,** RhoG-hepcidin binding to NW11-FPN in the presence of 10 μM $CoCl_2$ measured by fluorescence polarization reveals a $K_D$ of 7.7 nM. **d,** Fluorescence polarization of RhoG-hepcidin increases further with Fab45D8, consistent with formation of a larger complex. Importantly, Fab45D8 does not decrease RhoG-hepcidin binding. **e,** RhoG-hepcidin binds to FPN (FPN+hepcidin) but not to Fab45D8 alone (Fab45D8+hepcidin). Order of hepcidin and Fab45D8 addition does not influence the increase in fluorescence polarization for a Fab45D8-FPN-hepcidin complex.

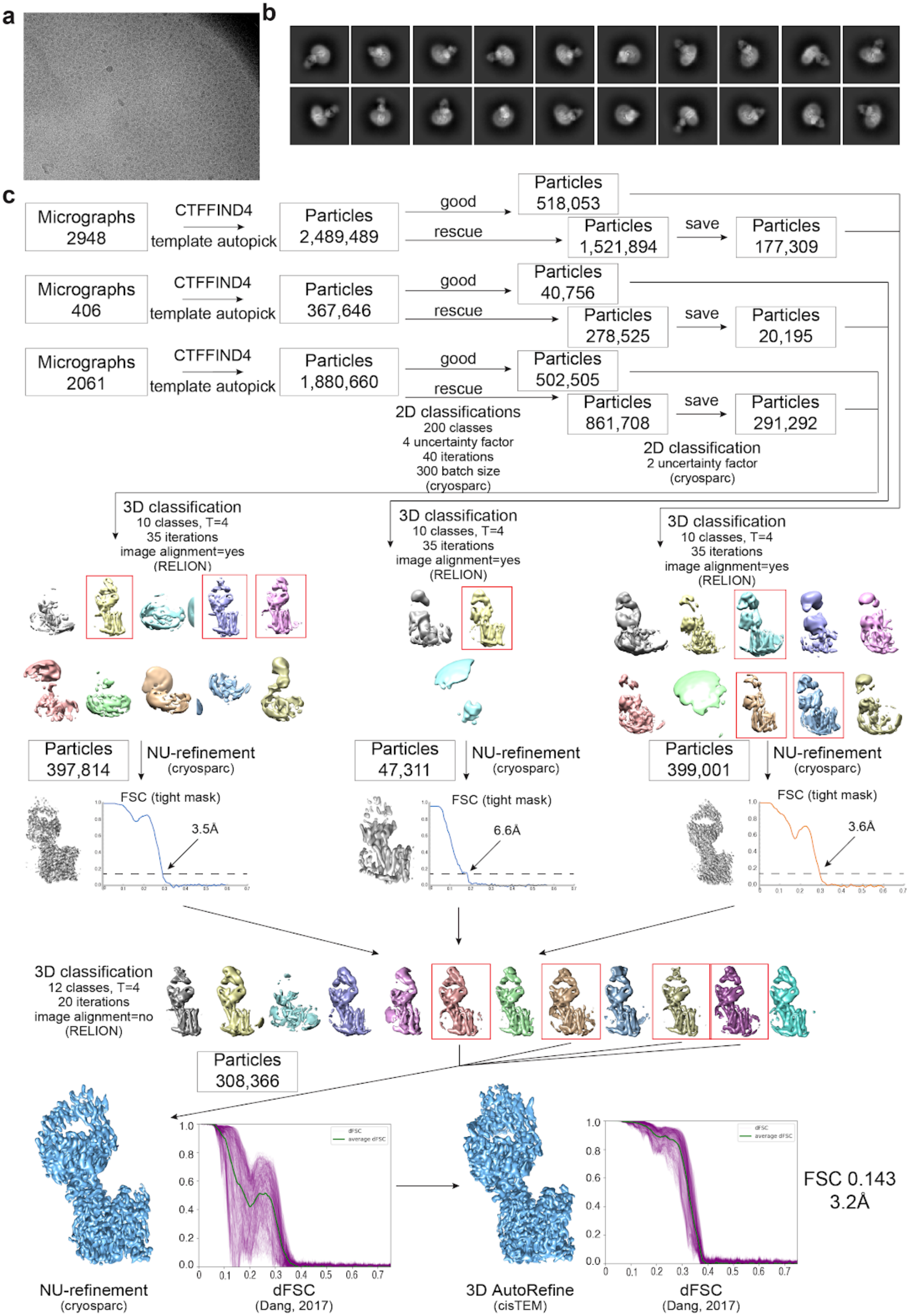
a
b
c
Micrographs 2948
CTFFIND4
template autopick
Particles 2,489,489
good
rescue
Particles 518,053
Particles 1,521,894
save
Particles 177,309
Micrographs 406
CTFFIND4
template autopick
Particles 367,646
good
rescue
Particles 40,756
Particles 278,525
save
Particles 20,195
Micrographs 2061
CTFFIND4
template autopick
Particles 1,880,660
good
rescue
Particles 502,505
Particles 861,708
save
Particles 291,292
2D classifications
200 classes
4 uncertainty factor
40 iterations
300 batch size
(cryosparc)
2D classification
2 uncertainty factor
(cryosparc)
3D classification
10 classes, T=4
35 iterations
image alignment=yes
(RELION)
3D classification
10 classes, T=4
35 iterations
image alignment=yes
(RELION)
3D classification
10 classes, T=4
35 iterations
image alignment=yes
(RELION)
Particles 397,814
NU-refinement
(cryosparc)
Particles 47,311
NU-refinement
(cryosparc)
Particles 399,001
NU-refinement
(cryosparc)
FSC (tight mask)
3.5Å
FSC (tight mask)
6.6Å
FSC (tight mask)
3.6Å
3D classification
12 classes, T=4
20 iterations
image alignment=no
(RELION)
Particles 308,366
NU-refinement
(cryosparc)
dFSC
(Dang, 2017)
3D AutoRefine
(cisTEM)
dFSC
(Dang, 2017)
FSC 0.143
3.2Å

**Supplementary Figure 3. Cryo-EM data processing for FPN-Fab45D8. a,** Representative motion-corrected micrograph collected on the Titan Krios showing monodisperse FPN-Fab45D8 nanodisc particles. **b,** Examples of “good” 2D class averages that were used in 3D classification. **c,** Flowchart showing image processing pipeline for FPN-Fab45D8. Initial processing, through 2D classification, was performed in cryoSPARC. Particles were then transferred, using csparc2star.py, to RELION for 3D classification, then to cryoSPARC for a nonuniform refinement, and finally to cisTEM for refinement. The number of particles moving into each step are noted. **d,** Final refinements from cryosparc and cisTEM beside their directional FSC curves calculated using dfsc.0.0.1.py.

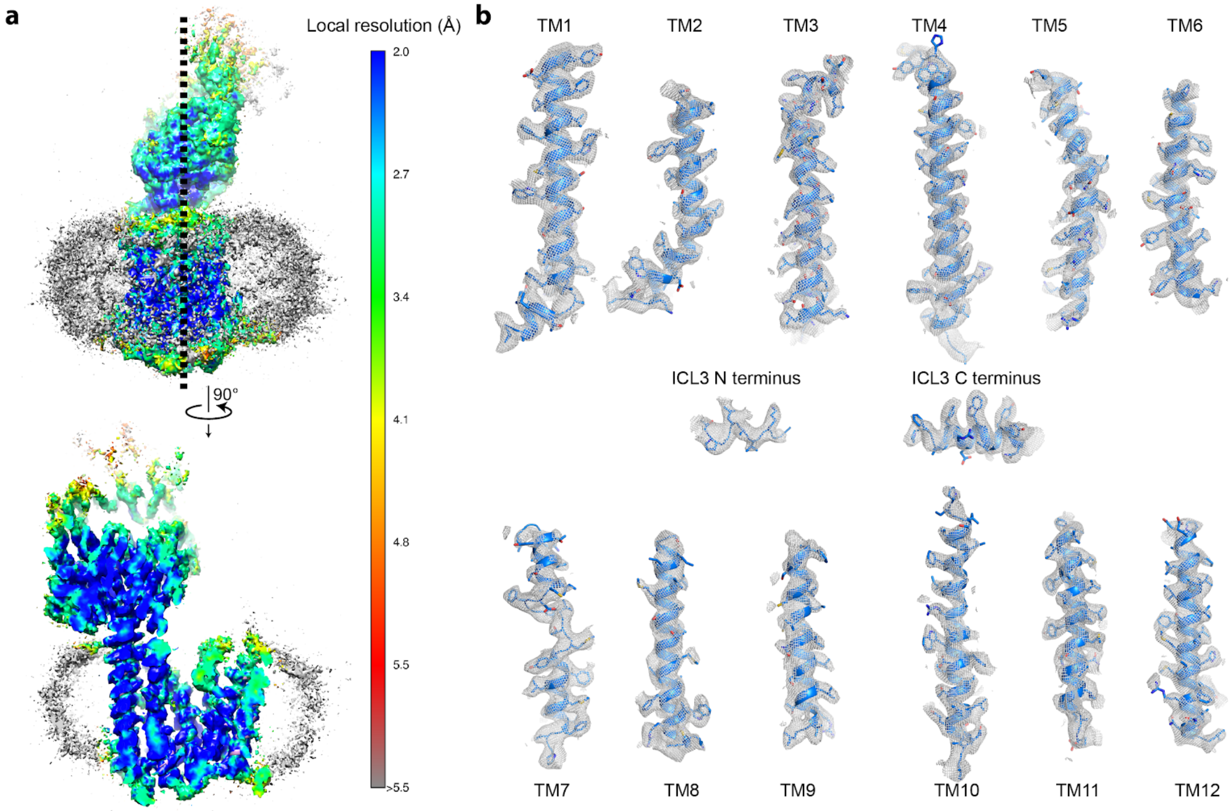


**Supplementary Figure 4. Cryo-EM map density for FPN. a,** Local resolution estimation by ResMap. Two views of the FPN-Fab45D8 complex in nanodisc are shown. The local resolution is highest for the FPN N domain **b,** Shown in grey mesh is cryo-EM map density for individual transmembrane helices. Mesh depicts density within a 2.5 Å radius of any modeled atom.

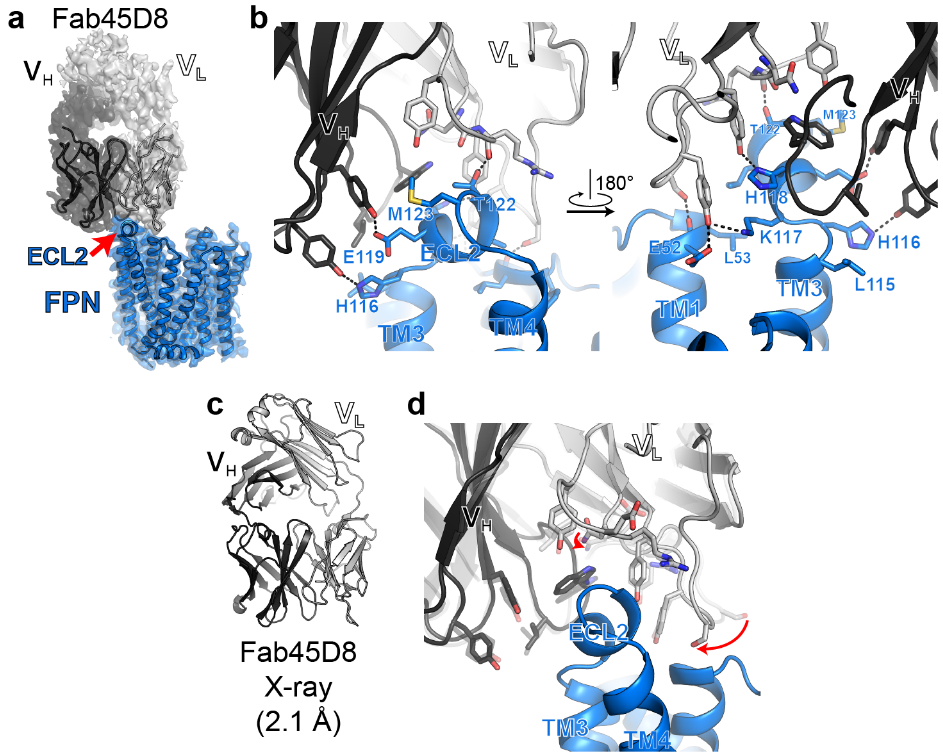


**Supplementary Figure 5. Structure of Fab45D8 and interaction with FPN. a,** Cryo-EM density of FPN and Fab45D8. Although density is observed for the constant regions of Fab45D8, it is not of sufficient quality to unambiguously model. **b,** Fab45D8 makes extensive contacts with FPN extracellular loop 2 (ECL2) with both the heavy ($V_H$) and light ($V_L$) chains. **c,** Crystal structure of Fab45D8 at 2.1 Å. **d,** Comparison of Fab45D8 alone (transparent cartoon and sticks) and bound to FPN. The binding site residues of Fab45D8 change minimally upon binding FPN.

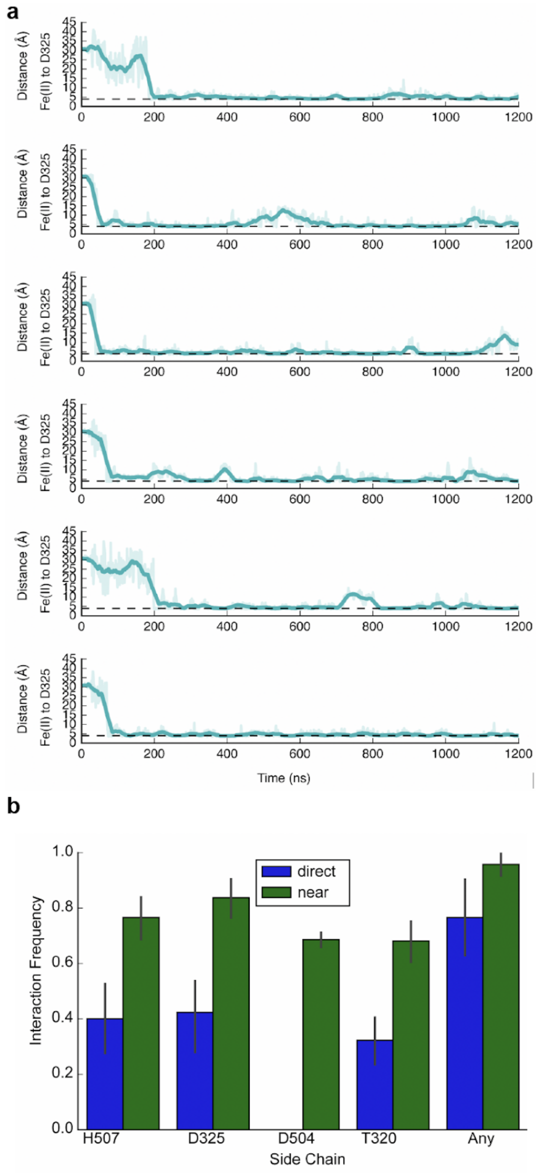


**Supplementary Figure 6. Molecular dynamics simulations of iron binding to FPN. a,** Each graph corresponds to an independent simulation where $Fe^{2+}$ ions start in solution and bind spontaneously to the proposed iron binding site. Distance shown is from the ion to the nearest oxygen atom of the D325 side chain. Thick traces represent a 15-ns sliding mean and thin traces represent unsmoothed values. Timetraces include 30 ns of equilibration. The average time to bind (the time from start of production simulation to when the measured distance is less ) was 71 ns. **b,** Using iron-bound simulations (a $Fe^{2+}$ ion placed in the proposed iron binding site), we quantified the fraction of simulations time that iron formed interactions with the side chains in the binding site. A direct coordinating interaction is designated here as when the distance between $Fe^{2+}$ and coordinating atom is <3 Å. A near interaction is characterized by a distance of <5.5 Å. “Any” describes the situation where any of the 4 preceding interactions are direct or near, respectively. Data is averaged from 6 independent simulations, error bars are s.e.m.

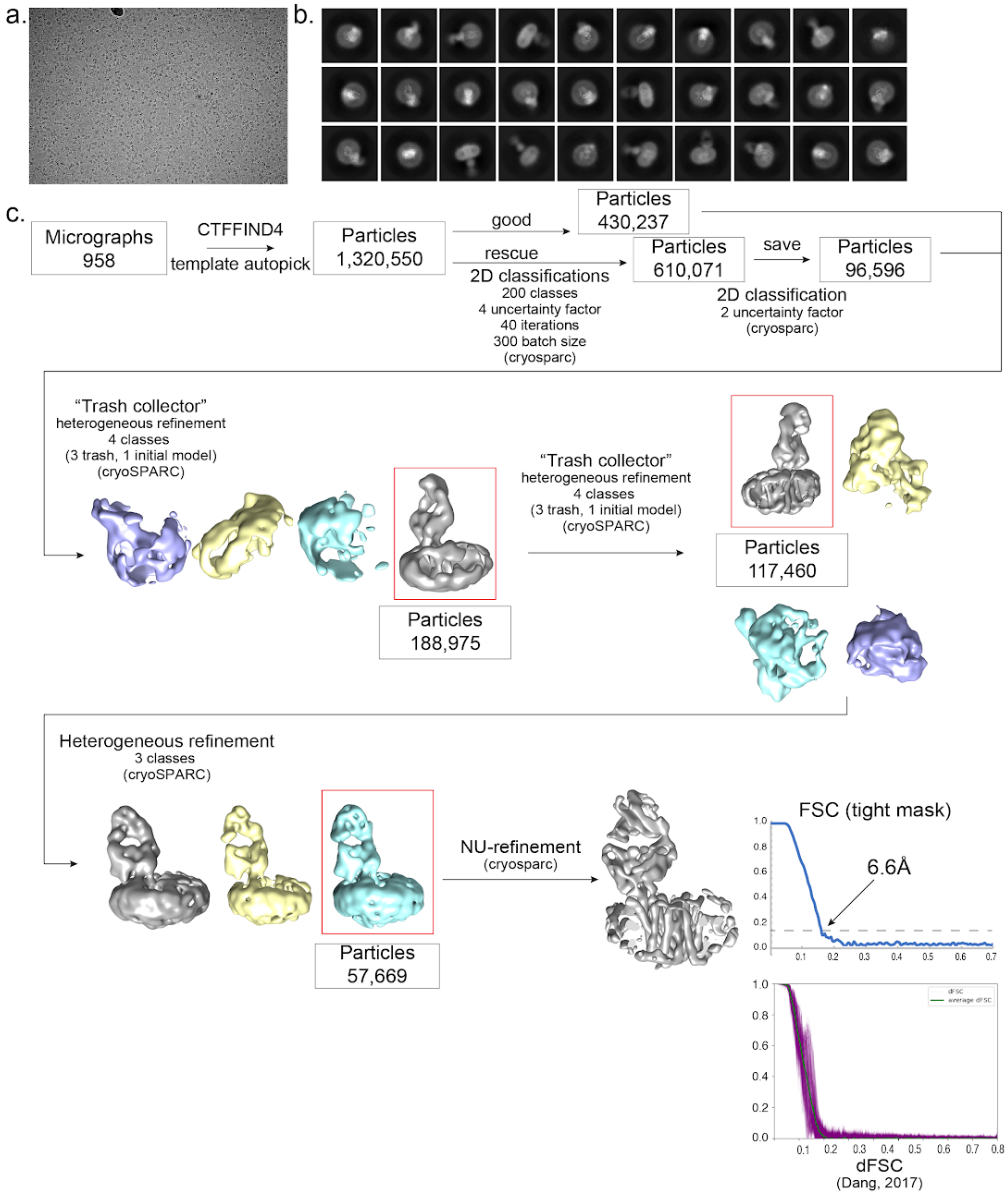


**Supplementary Figure 7. Cryo-EM data processing for FPN-Fab45D8-$Co^{2+}$-hepcidin. a,** Representative motion-corrected micrograph collected on the Talos Arctica operated at 200kV. showing monodisperse FPN-Fab45D8-$Co^{2+}$-hepcidin nanodisc particles. **b,** Examples of "good" 2D class averages that were sent into 3D classification. **c,** Flowchart showing image processing pipeline for hepcidin and $Co^{2+}$ bound FPN-Fab45D8. All processing was performed in cryoSPARC with details in the methods section. The number of particles moving into each step are noted. A final non-uniform refinement was performed in cryosparc the directional FSC curve was calculated using dfsc.0.0.1.py.

**Supplementary Table 1: Cryo-EM statistics**

| | FPN-Fab45D8 (EMDB-21539) (PDB 6W4S) | FPN-$Co^{2+}$-hepcidin-Fab45D8 (EMDB-21550) |
|---|---|---|
| **Data collection and processing** | | |
| Microscope/Detector | Titan Krios/Gatan K3 with Gatan Bioquantum Energy Filter | Talos Arctica/Gatan K3 |
| Imaging software and collection | SerialEM, 3x3 image shift | Serial EM, 3x3 image shift |
| Magnification | 105,000 | 28,000 |
| Voltage (kV) | 300 | 200 |
| Electron exposure (e–/Å$^2$) | 66 | 60 |
| Dose rate (e–/pix/sec) | 8 | 10 |
| Frame exposure (e–/Å$^2$) | 0.55 | 0.5 |
| Defocus range (µm) | -0.8 to -2.0 | -1.0 to -2.0 |
| Pixel size (Å) | 0.834 (physical) | 0.7 (physical) |
| Micrographs | 5,415 | 958 |
| **Reconstruction** | | |
| Autopicked particules (template-based in cryosparc) | 4,737,795 | 1,320,550 |
| Particles in 3D classification (RELION) | 1,326,130 | 526,833 |
| Particles in final refinement | 308,386 (cisTEM) | 57,669 (cryosparc) |
| Symmetry imposed | C1 | C1 |
| Map sharpening *B* factor (Å$^2$) | -130 | -600 |
| Map resolution, global FSC (Å) | | |
| FSC 0.5, unmasked/masked | 7.1/3.6 | 12.5/8.5 |
| FSC 0.143, unmasked/masked | 4.0/3.2 | 8.0/6.5 |
| **Refinement** | | |
| Initial model used (PDB code) | 5AYN | |
| Model resolution (Å) | | |
| FSC 0.5, unmasked/masked | 3.6/3.4 | |
| Model composition | | |
| Non-hydrogen atoms | 4968 | |
| Protein residues | 650 | |
| *B* factors (Å$^2$) | | |
| Protein | 67.31 | |
| R.m.s. deviations | | |
| Bond lengths (Å) | 0.004 | |
| Bond angles (°) | 0.563 | |
| Validation | | |
| MolProbity score | 2.31 | |
| Clashscore | 7.90 | |
| Poor rotamers (%) | 4.45 | |
| EMRinger score | 2.37 | |
| CaBLAM score | 2.06 | |
| Ramachandran plot | | |
| Favored (%) | 94.38 | |
| Allowed (%) | 5.62 | |
| Disallowed (%) | 0 | |

**Supplementary Table 2: Fab45D8 crystal structure statistics**

| | Fab45D8 (PDB 6W4V) |
|---|---|
| **Data collection** | |
| Space group | $P2_1$ |
| Cell dimensions | |
| $a$, $b$, $c$ (Å) | 72.19, 36.43, 84.95 |
| α, β, γ (°) | 90.0, 112.26, 90.0 |
| Resolution (Å) | 39.31 - 2.09 (2.14 - 2.09)[a] |
| $R_{sym}$ or $R_{merge}$ | 0.087 (1.385) |
| $I / \sigma I$ | 7.9 (1.2) |
| Completeness (%) | 99.9 (99.9) |
| Redundancy | 3.3 (3.5) |
| CC (1/2) (%) | 99.6 (30.0) |
| **Refinement** | |
| Resolution (Å) | 39.31 - 2.09 |
| No. reflections | 24736 |
| $R_{work}$ / $R_{free}$ (%) | 21.6 / 24.7 |
| No. atoms | |
| Protein | 3356 |
| Ligand/ion | 20 |
| Water | 125 |
| $B$-factors | |
| Protein | 57.87 |
| Ligand/ion | 86.17 |
| Water | 51.43 |
| R.m.s. deviations | |
| Bond lengths (Å) | 0.01 |
| Bond angles (°) | 0.92 |

[a]Values in parentheses are for highest-resolution shell.

**Supplementary Table 3: Molecular dynamics simulations**

| Condition | Number of Simulations | Duration (μs) | Notes |
|---|---|---|---|
| Iron absent | 6 | 2.2 (each) | No iron was added |
| Iron bound | 6 | 2.2 (each) | A single $Fe^{2+}$ ion was placed in the proposed binding site initially |
| Iron in bulk solvent | 6 | 2.2 (each) | 15 $Fe^{2+}$ ions were placed randomly in solution initially |
